# Robust Phylogenetic Tree-based Microbiome Association Test using Repeatedly Measured Data for Composition Bias

**DOI:** 10.1101/2023.07.10.548346

**Authors:** Kangjin Kim, Sungho Won

## Abstract

**Motivation:** The effects of microbiota on the host phenotypes can substantially differ depending on his/her age. Longitudinally measured microbiome data allows us to detect the age modification effect and are useful for the detection of microorganisms related to the progression of disease which change identification over time. Moreover, longitudinal analysis enables the estimation of within-subject covariate effect, is robust against the between-subject confounders, and provides better evidence for the causal relationship than cross-sectional studies. However, they suffer from compositional bias, and few statistical methods can estimate their effect on host diseases with repeatedly measured 16S rRNA gene data. In this article, we proposed mTMAT which can be applied to longitudinal microbiome data and is robust against compositional bias.

**Results:** mTMAT normalized the microbial abundance and utilized the ratio of the pooled abundances for association analysis. mTMAT is based on generalized estimating equations with a robust variance estimator and can be applied to repeatedly measured microbiome data. With extensive simulation studies, we showed that mTMAT is statistically more powerful and is robust against compositional bias. mTMAT enables detection of microbial taxa associated with host diseases using repeatedly measured 16S rRNA gene data and can provide deeper insight into bacterial pathology.

**Availability:** The 16S rRNA amplicon sequencing metagenomics datasets for Korea Association REsource cohort is available from the NCBI Sequence Read Archive database under project accession number PRJNA716550. mTMAT was implemented in the R package. Detailed information is available at https://healthstat.snu.ac.kr/software/mtmat.

**Supplementary information:** Supplementary data are available at *Bioinformatics* online.

## Introduction

Recent advancement in high-throughput technologies such as microarrays and next-generation sequencing has significantly elucidated the microbial world. For instance, intestinal microbiota has been found to play essential roles in the host by influencing energy homeostasis, body adiposity, blood sugar control, insulin sensitivity, and hormone secretion [1-3]. However, the composition of the microbiota varies from subject to subject, and the abundances of microbial taxa are often sparse with excessive zeros. Furthermore, the microbiota is affected by various factors, such as age and sex, which cause its abundance to vary considerably. Such sparsity in the data makes controlling type-1 and type-2 errors in statistical analyses difficult, and the inference of causal relationships through statistical analysis of microbiota data should be performed cautiously.

Repeatedly measured microbiota studies are useful for the detection of microorganisms related to the progression of disease and change identification over time, and provide more evidence for the causal relationship than cross-sectional studies [4]. Furthermore, the estimation of within-subject covariate effects is robust against between-subject confounders, and repeatedly measured microbiome data allow for the robust identification of microbiota effects on the risk of diseases in the host. Statistical analyses with repeatedly observed 16S rRNA gene require the adjustment of similarity among the measurements of the same subjects. However, only a limited number of methods are applicable in the repeated observation of 16S rRNA gene data, and the development of statistical methods for longitudinal studies is required for investigating the association between the human microbiome and diseases.

Xia et al [5] comprehensively reviewed statistical methods of longitudinal data analysis in microbiome studies. These methods can be categorized into several categories: (1) standard longitudinal model, (2) overdispersed and zero-inflated longitudinal models, and (3) multivariate distance/kernel-based longitudinal models. First, the standard longitudinal model includes the linear mixed effect model (LMM) with generalized estimation equation (GEE) and generalized linear mixed effect model (GLMM) among others. LMM provides a standardized and flexible approach toward modeling both fixed and random effects. However, operational taxonomic unit (OTU) abundances cannot address the sparsity issue and should be transformed or normalized to avoid the violation of distribution assumptions. Second, overdispersed and zero-inflated longitudinal models include the zero-inflated Gaussian (ZIG) mixture model, extensions of negative binomial mixed-effects (NBMM) [6], and zero-inflated negative binomial models (FZINBMM) [7]. The two-part zero-inflated beta regression model with random effects (ZIBR) extends the zero-inflated beta regression model to longitudinal data settings [8]. FZINBMM and ZIBR can analyze overdispersed and zero-inflated longitudinal metagenomics data. Finally, the multivariate distance/kernel-based longitudinal model includes the correlated sequence kernel association test (cSKAT) for continuous outcome, and the generalized linear mixed model and its data-driven adaptive test (GLMM-MiRKAT) for non-normally distributed outcome such as binary traits.

However, in spite of such developments, longitudinal microbiome data suffers from compositional bias, and only a few methods are robust against this problem. The magnitude of the sequence depth differs from subject to subject in metagenomics data, and the sums of absolute abundances for each subject are substantially different; therefore, relative abundances are generally used. However, this generates compositional bias, especially in longitudinally observed microbiome data [9], because relative abundances at each time point only provide a proportion of a taxa and it is therefore not possible to compare the relative abundance of the same subject. Furthermore, relative abundances are correlated among different taxa, and their adjustment is necessary if they were utilized as a response variable [10].

Several methods have been proposed to adjust the compositional bias. For instance, additive log-ratio (ALR) that uses a reference abundance for its denominator and centered log-ratio (CLR) that uses geometric mean can be considered. Network analysis, including sparse correlations for compositional data and sparse inverse covariance estimation for ecological association inference, can also be considered, modeling the whole community with a statistical model. However, they can only be applied to cross-sectional data, and none of them can be applied to analyzing longitudinally observed datasets [11, 12].

In this article, we propose the phylogenetic tree-based microbiome association test for repeatedly measured data (mTMAT), which pools the abundance of OTUs based on the phylogenetic distance, thus correcting the problem of zero-inflation, making it robust against compositional bias. Through extensive simulation and real data analyses, we prove its robustness against compositional bias and its improved statistical power compared to other methods.

## Methods

### 2.1 Phylogenetic tree

The same notations and assumptions as those of TMAT were applied [13]. We denoted the absolute abundance of OTU *m* of subject *i* at time point *j* to be 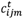, where *i =* 1, …, *N, j =* 1, …, *N*_*i*_, *m* = 1, …, *M*. We assumed that OTUs are clustered by profiling the sequences of all subjects at all the time points simultaneously, and a rooted binary phylogenetic tree was provided for these OTUs. The first *M*_1_ OTUs belong to a taxonomy of interest *t*, where *t* = 1, … *T*, for the analysis of its association with host diseases, while the other *M* – *M*_t_ OTUs belong to a different taxonomy. The genus with the first *M*_1_ OTUs has *M*_t_ – 1 internal nodes and *M*_1_ leaf nodes. Internal nodes are denoted by *k*, where *k* =1, …, *M*_t_ – 1. Leaf nodes are denoted by *m*, where *m* = 1, …, *M*. For each leaf node, there is a corresponding single OTU; if *m* = 1, …, or *M*_t_, *m* is the leaf node of the genus of interest, otherwise *m* belongs to a different genus. The absolute abundance of OTU *m* of subject *i* at time point *j* is denoted by 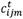. Assuming that mutations are transmitted from the left (right) child node to all of its leaf nodes, the relative proportion of the leaf nodes of the left child node increases for cases where the mutation occurs during transmission to its left child node and decreases if it does so during transmission to the right child node. When testing the association of an internal node *k* with a host disease of interest, the internal node *k* and its leaf nodes are considered to be representative of a test node and test leaf nodes, respectively. The left and right test leaf nodes further represent the leaf nodes of the left and right child nodes of a test node, respectively.

For internal node *k* in the genus with *M*_t_ OTUs, *L*_*k*_ and *R*_*k*_ are regarded as the sets of its left and right leaf nodes, respectively. Figure 1 illustrates these definitions.

**Figure 1.**
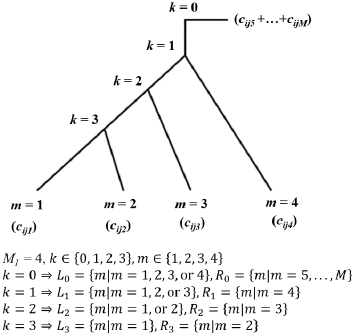
Examples of rooted binary phylogenetic trees.

### 2.2 Quasi-likelihood

The log-transformed CPM *r*_*ijm*_, which is used for the edgeR package (version 3.16.5) is expressed as follows.

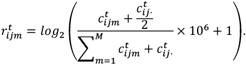

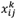, where *k* = 1, …, *M*_1_ – 1, is expressed as follows

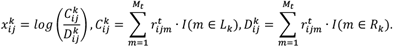

Here I(A) indicates the indicator function. As all OTUs in taxonomy *t* can be associated with the host disease, 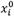 for such a case is expressed as follows

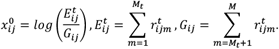

Here, we assume that ***x***^0^, …, and 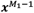 are ordered according to the depth of the internal nodes. ***x***^0^ is used to test the association of all OTUs belonging to the genus of interest by pooling them, and ***x***0 is used for testing the root node of the phylogenetic tree. The phenotype of subject *i* at time point *j* is denoted by *y*_*ij*_ and is coded as 1 and 0 for cases and controls, respectively. Two additional options for the value of *G*_*ij*_ are provided to consider compositional bias. The first one uses 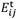 value of a reference taxon while the second one uses the geometric mean 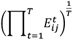 for all taxonomies. These two options correspond to the ALR and CLR transformations, respectively. The vectors and matrices for testing the association of the genus of interest are expressed as follows.

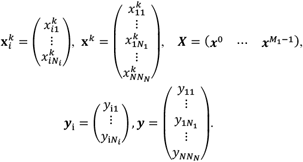

Based on such matrix notations, we provide the quasi-likelihood for repeatedly observed 16S rRNA gene data. If we denote ***R***_*i*_ and *σ*_*kk*_ as a working correlation matrix and overdispersion parameter, respectively, and define ***D***_*ik*_ as a diagonal matrix with its diagonal entries being 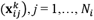, the covariance matrix for the observations of subject *i* is expressed as follows

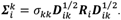

Then the covariance matrix **Σ**^*k*^ can be expressed as follows

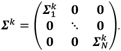

If we let Z be a design matrix for *p* covariates, including the intercept, we assume

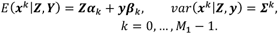

Therefore, quasi-score functions for ***α***_***k***_ and ***β***_*k*_ can be expressed as follows

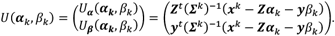

Quasi-fisher information can be expressed as follows

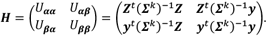

### 2.3 Score test with small sample adjustment

The null hypothesis can be expressed as follows

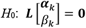

where ***L*** is a matrix of linear constraints with *c* rows and number of columns equal to the length of 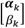 and **0** is the zero vector of matching dimension. To test the null hypothesis *H*_0_: *β*_*k*_ = 0, the generalized score statistics by Boos can be applied [14] by setting ***L*** = [**0**^*t*^ 1] as follows:

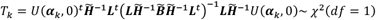

where

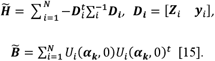

To adjust the small sample bias, 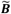 is further updated as follows

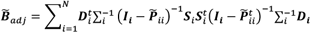

where

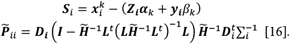

### 2.4 Wald test with small sample adjustment

The Wald statistic with sandwich estimator with correction of small sample bias was considered [15]. ***β***_*k*_ can be estimated by solving the estimating equation, *U*_***β***_(***α***_***k***_, ***β***_*k*_) = 0, as follows

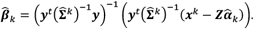

For the estimation of the variance of 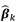 we consider the robust variance estimator with a small sample adjustment as follows

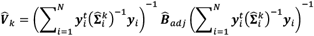

where

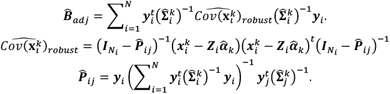

Based on this result, the robust Wald statistic of ***β***_*k*_ for test node *k* is expressed as follows

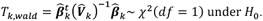

### 2.5 mTMAT

Statistics for *H*_0_: *β*_*k*_ = 0 can be combined to test 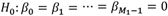 using the minimum p-value. If p-values for *T*_*k*_ are denoted by *pT*_*k*_, the proposed statistics, mTMAT_M_, are expressed as follows

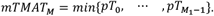

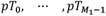 are asymptotically independent [13]. Therefore, we can conclude that

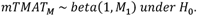

If the sample size is small, the normality of *T*_*k*_ under *H*_0_ may not be achieved, and the assumption of the quasi-score test can be violated. If we apply the inverse normal transformation to 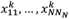, then the same statistics can be obtained. This is denoted by 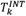. Rank-based inverse normal transformation with an adjustment parameter of 0.5 was used for the transformation, and data with tie values were mapped to the same value in the transformed data [17]. Then, mTMAT_IM_ is expressed as follows

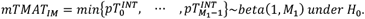

### 2.6 KARE Cohort data

The Korea Association REsource (KARE) cohort is a prospective study cohort involving subjects from the rural community of Ansung and the urban community of Ansan in South Korea. It began in 2001 as part of the Korean Genome Epidemiology study [18]; we used data from 2,072 urine samples from 691 participants in 2013, 2015, and 2017. Their 16S rRNA gene amplicon sequencing data used in the study were obtained from the NCBI Sequence Read Archive database under project accession number PRJNA716550. For paired-end sequencing of the V3-V4 region of the bacterial 16S rRNA gene, the widely used primers 16S_V3_F (5’-TCGTCGGCAGCGTCAGATGTGTATAAGAGACAGCCTAC-GGGNGGCWG CAG-3’) and 16S_V4_R (5’-GTCTCGTGGGCTCGGAGATGTGTAT-AAGAGA CAGGACTACHVGGGTATCTAATCC-3’) were used. Adaptor sequences were detected and removed using the CUTADAPT software with a minimum overlap of 11 bp, maximum error rate of 10%, and a minimum length of 10 bp [19]. Sequences were merged using CASPER with a mismatch ratio of 0.27 and filtered by the Phred (Q) score, resulting in sequences of 350–550 bp in length [20, 21]. After the merged sequences were dereplicated, chimeric sequences were detected and removed using VSEARCH and the Silva Gold reference database for chimeras [22]. The open-reference Operational Taxonomic Unit (OTU) picking was based on the EzTaxon database using UCLUST [23, 24]. The phylogenetic trees based on EzTaxon database were obtained through the SINA method [25] using the reference sequences available from the EzTaxon database. We calculated the proportion of each OTU among the total OTUs and determined the mean value across all subjects. If the resulting value was <0.001, the OTU was excluded [26]. Among the 691 subjects, those with a read count of <3,000 or for whom genomic data were not available in any phase were excluded. As a result, 1179 samples from 393 subjects, including 70 genera, were used for the simulation analysis.

### 2.7 Simulation studies

We conducted extensive simulations to evaluate the performance of mTMAT with two types of datasets. One dataset with 393 subjects who participated in all three phases from KARE cohort and the generative dataset based on microbiomeDASim [27]. The disease status of the subjects was permuted. A single test node was randomly selected from the internal nodes, and from their test leaf nodes, a single OTU, 50% of OTUs, or 90% of OTUs was randomly selected as causal OTUs. These were denoted by p = 1 OTU, 50%, and 90%, respectively. p = 1 indicates a single OTU associated with the host disease; therefore, the phylogenetic tree structure does not provide any useful information for mTMAT. If we let the sample variances of *c*_*im*_ for causal OTUs be 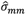, the observed absolute abundances of the selected causal OTUs for affected subjects was assumed to be 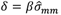, where *β* = 0, 0.01, 0.02, or 0.04, and the absolute abundances of the other OTUs were used without modification. *β* = 0 was considered for estimation of empirical type-1 error rates, and the others were used for estimating statistical power. Type-1 error rates were estimated at the 0.1, 0.05, 0.01, and 0.005 significance levels with 5,000 replicates. Empirical power was estimated at the 0.05 significance level with 500 replicates.

For comparison with mTMAT_M_ and mTMAT_IM_, GLMM-MiRKAT (version 1.2), FZINBMM (version 1.0), linear mixed model (LMM) with arcsine square root transformation (LMM-arcsine), and LMM with log transformation (LMM-log) with nlme package (version 3.1) were considered. Phylogenetic tree-based microbiome association test (TMAT, version 1.01), optimal microbiome regression-based kernel association test (oMiRKAT, version 0.02), adaptive microbiome-based sum of powered score (aMiSPU, version 1.0), and the Wilcoxon test were also considered for comparison with cross-sectional methods. Association analyses were conducted at the genus level. FZINBMM, LMM models, and Wilcoxon were applied by pooling all OTUs within each genus. Each genus consisted of multiple OTUs, and oMiRKAT and aMiSPU were applied to OTUs belonging to each genus.

For mTMAT_M_ and mTMAT_IM_, robust wald and score statistics with four different choices of working correlation matrix: identity, compound symmetry (CS), first-order autoregressive (AR1) and unstructured (UN), were considered. aMiSPU and oMiRKAT used permutation-based p-values and were calculated with 500 and 5,000 permutated replicates for estimation of power and type-1 error rates, respectively. GLMM-MiRKAT and oMiRKAT offer several distance metrics, including Unifrac distance as a default choice, while aMiSPU also uses Unifrac distance as the default option; we considered the default choices. However, Unifrac distance cannot be calculated if read counts are not observed. Therefore, subjects with no read counts were excluded from GLMM-MiRKAT, oMiRKAT, and aMiSPU. Furthermore, none of these can analyze a genus with a single OTU; therefore, such instances were not considered for statistical power estimations of such genera.

Following the simulation with the generated dataset with microbiomeDASim, Identity, CS, and AR1 with different parameter values are assumed for the simulation, and type-1 error estimates were compared for different uses of working correlation matrices for mTMAT_IM_. The mean value of relative abundance and proportion of zero count samples were estimated from the KARE cohort study for all the genera; the genera with first quantile, median, and third quantile sparsity level were selected for the simulation. The values were 52%, 64%, and 73%.

We also evaluate the robustness of the proposed method against the compositional bias. Different choices of *G*_*ij*_ corresponds to ALR and CLR are also considered. The KARE dataset was simulated 2000 times with simulation parameters *N* and the ratios of the cases and controls were 50 and 1:3, respectively. Then a genus containing more than one OTU was selected and assumed to be associated with a phenotype with *β* = 0.15 and *p* =50%. Then an OTU contained in the chosen genus was selected and set to be associated with a phenotype with the same *β*. The abundance of the selected OTU that is not in the chosen genus was added by its standard deviation multiplied by multipliers 0, 1, 5, 10, 50, and 100. Then the mean and interquartile range of bias estimate of the selected genus was calculated and compared with a different value of the multiplier.

### 2.8 Pregnant microbiome data

We used publicly available datasets from Romero [28], who conducted a retrospective case–control longitudinal study that included non-pregnant women (n = 32) and pregnant women who delivered at term (38 to 42 weeks) without complications (n = 22) using pplacer and version 0.2 of the vaginal community 16S rRNA gene database [29] for the taxonomic classification, and the neighbor joining method based on the Bray-Curtis dissimilarity index is used to obtain phylogenetic tree [30]. The pregnant dataset includes data on the race, days after the first visit (GDColl), household income, maternal education, and baby gender.

## Results

### 3.1. Results from simulated data

The performances of mTMAT_M_ and mTMAT_IM_ were evaluated using simulated data. Supplementary Figure 1 shows the overall distribution of microbial composition. Supplementary Table 1 shows that inflation of type-1 error was observed when the case samples and total samples increased. There appears to be no notable difference of type-1 error rates for the choice of working correlation matrix. Conversely, mTMAT_IM_ adequately preserved the nominal type-1 error with a slight inflation when unstructured correlation (Table 1).

**Table 1.**
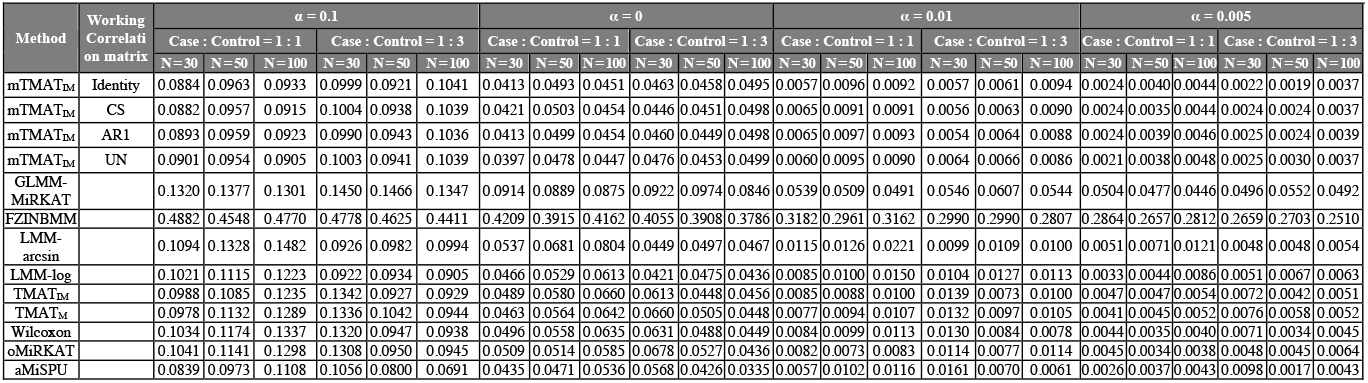
Type-1 error estimates of mTMATIM and other statistical methods from repeatedly measured microbiome data at 3 time points. The ratios between cases and controls were assumed to be 1:1 and 1:3. The total sample size is denoted by N, and we considered N = 30, 50, and 100. All subjects were selected without replacement. Type-1 error estimates were calculated with 2,000 replicates at the significance levels 0.1, 0.05, 0.01 and 0.005.

GLMM-MiRKAT, FZINBMM and LMM models are designed to be used as longitudinal microbiome data and can be compared with mTMAT_IM_ and mTMAT_M_. FZINBMM and GLMM-MiRKAT could not preserve type-1 error rates with extremely high type-1 error estimates for FZINBMM. GLMM-MiRKAT suffered singular matrix problem during calculating the test statistics (Table 1). In this case, the resulting p-values were excluded for the estimation of type-1 error rate and power.

We also evaluated the effect of the number of leaf nodes (Supplementary Table 2), and the results showed that mTMAT_M_ became slightly conservative when the number of leaf nodes exceeded 5, but that mTMAT_IM_ was less affected. The result with more than 15 leaf nodes can be dependent on specific genera chosen with a small number of genera.

Supplementary Table 3 shows the effect of sparsity on the type-1 error rate. We calculated the sparsity of each genus, defined as the proportion of subjects with no abundance, and type-1 error rates were calculated. Results showed that the type-1 error rates of FZINBMM were the most inflated and that some inflation was observed for GLMM-MiRKAT when the mean sparsity exceeded 20%. GLMM-MiRKAT is based on the permutation, and the permutation-based p-value is generally robust to the non-normality. However, if heteroscedasticity exists, its statistical validity can be impaired. A substantial amount of sparsity may induce heteroscedasticity, which may explain the type-1 error inflation. Some inflation was also observed for TMAT_M_, but TMAT_IM_ rates were well preserved.

The effect of the assumed correlation matrix for different scenarios was evaluated with the use of microbiomeDASim [27]. Identity, CS, and AR1 with different parameter values are assumed for the simulation with different uses of working correlation matrix, robust score statistic and for mTMAT_IM_ (Supplementary Table 4). The result shows that mTMAT_IM_ preserved type-1 error for all the scenarios.

We also calculated statistical power estimates with 2,000 replicates at the 0.05 significance level, and compared them with those of other statistical methods. The significance levels for each method were adjusted based on the statistics from the simulation to calculate type-1 error for a valid performance comparison. The threshold is determined as the percentiles of the p-values calculated in the type-1 error simulation under null hypothesis. We considered genera comprising two or more OTUs. In Figure 2, mTMAT_IM_ usually outperformed the other methods. The performance of GLMM-MiRKAT was comparable with mTMAT_M_. FZINBMM and LMM-log exhibited significantly less power than other methods.

**Figure 2.**
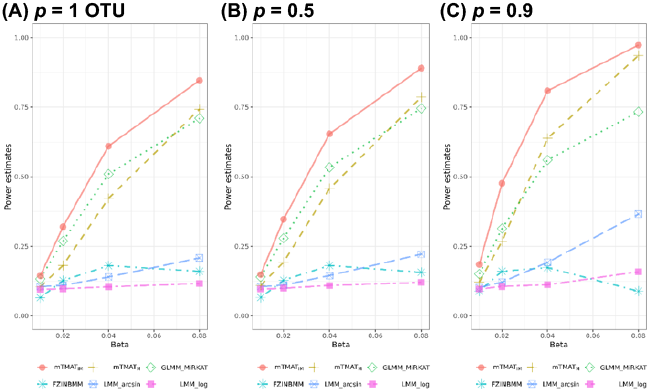
Power estimates for genera consisting of more than one OTU. Power estimates at the significance level 0.05 were calculated with 500 replicates. We generated simulation data based on read counts from datasets, and considered genera with more than one OTU. For statistical methods whose type-1 errors are violated, their P-values were adjusted with the simulated data, which makes all statistical methods preserve the nominal significance level. We assumed the total sample size (N) is equal to 50 and the ratio of cases and controls is set to be 1:3. Identity working correlation matrix and robust score statistic are used for mTMAT.

Supplementary Figure 2 shows the results of genera comprising one or more OTUs. GLMM-MiRKAT can only be calculated if more than one OTU is available. Therefore, it was excluded from this comparison. mTMAT_M_ and mTMAT_IM_ can be applied in such scenarios, and the results showed that the proposed method was the most efficient.

Comparison with methods for cross-sectional analysis (Supplementary Figure 3) shows that TMAT_M_, TMAT_IM_, and mTMAT_IM_ showed high statistical power. aMiSPU had the highest power estimate when beta was 0.02.

Supplementary Figure 4 shows the statistical power estimates according to the number of leaf nodes. The number of causal OTUs was assumed to be the same, and statistical power estimates decreased according to the number of leaf nodes. Optimal performance was observed for mTMAT_IM_; however, GLMM-MiRKAT exhibited a high level of type-1 error rate with the leaf node ranging between 6 and 15. The sparsity levels were also considered, and statistical power estimates were compared. For each genus, sparsity was defined by the proportion of subjects with no reads from that genus. Power estimates were maximized at the middle-group sparsity level (Supplementary Figure 5). The type-1 error and the power of group and time and interaction effect are also estimated using simulation dataset with microbiomeDASim package. (Supplementary Table 5 and Supplementary Table 6). The nominal type-1 error of the group, time (Supplementary Table 5) and interaction effect (Supplementary Table 6) are well reserved for all the methods except GLMM-MiRKAT. We verified that mTMAT_IM_ and mTMAT_M_ can successfully capture group and time effects having the other as covariate. mTMAT_IM_ and mTMAT_M_ also detected the interaction effect. LMM-arcsine and LMM-log had shown higher power than mTMAT_IM_ and mTMAT_M_ for this simulation scenario. Supplementary Figure 6 shows the effect of compositional bias. mTMAT_IM_ and FZINBMM had a smaller bias than other methods when the level of compositional bias was high. All three types of mTMAT_IM_ had smaller interquartile ranges of bias than FZINBMM, indicating that mTMAT_IM_ successfully mitigated compositional bias compared to other methods.

In summary, we confirmed that mTMAT_IM_ is generally the most efficient method among those available in our simulations. mTMAT_IM_ considers phylogenetic tree structures, uses log CPM transformation, correction of compositional bias with taking a proportion among OTUs, and consider correlations among repeatedly measured samples, which makes it superior to other methods. The overall power comparison results for cross-sectional methods are consistent among previous studies on TMAT [13]; however, the type-1 error rate for TMAT was inflated with longitudinal microbiome data. GLMM-MiRKAT is the second most powerful but failed to preserve type-1 rates and cannot be applied in analysis with single OTU. Furthermore, GLMM-MiRKAT is based on oMiRKAT, and they both used the kernel method and permutation approaches, which can be very computationally intensive, especially if the sample size increases [13].

### 3.2. Real data analysis

The pregnant datasets were analyzed with mTMAT, GLMM-MiRKAT, FZINBMM, and LMM with the arcsine square root transformation and LMM with log transformation. The pregnant dataset includes the race, days after the first visit (GDColl), house hold income, maternal education, and gender of baby. The overall composition is presented in Supplementary Figure 7 and the overall composition change was clear after >300 days. Supplementary Figure 8 shows that the change can be related to the pregnancy state. PERMANOVA analysis result shows the associated phenotype that explained microbiome variability. Race was the most significant covariate, with a p-value of 0.06 (Supplementary

Figure 9). Table 2 shows that mTMAT_IM_ found 11 significant genera. FZINBMM, LMM-arcsine, and LMM-log found 16, 14, and 14 significant genera, respectively. As shown in the simulation study, most of the detected genera as significant only by FZINBMM can be false positives. mTMAT_IM_ shared most of the significant genera with other methods. The most significant genera was *Lactobacillus*, which is consistent with the findings of the original study [28]. Supplementary Figure 10 shows a Venn diagram comparing the numbers of significant genera implicated by the various applied methods. As LMM-arcsine and LMM-log differ only in their transformation, the methods shared all 16 detected genera. FZINBMM detected two more genera that were not detected by other methods. mTMAT_IM_ shared all the 12 detected genera with other methods.

**Table 2.** Association analysis results of Pregnant dataset. Results for genera significantly associated with at least one method at the FDR-adjusted 0.05 significance level were summarized.

| Family Taxonomy | Genus Taxonomy | mTMAT<sub>IM</sub> | mTMAT<sub>M</sub> | GLMM-MiRKAT | FZINBMM | LMM arcsine | LMM log |
| --- | --- | --- | --- | --- | --- | --- | --- |
| Lactobacillaceae | Lactobacillus | 0.01984 | 0.01549 | 0.00416 | 0.00000 | 0.00000 | 0.00000 |
| Tissierellaceae | Anaerococcus | 0.02110 | 0.02275 | 0.01247 | 0.00000 | 0.00000 | 0.00000 |
| Tissierellaceae | Peptoniphilus | 0.02599 | 0.02110 | 0.00416 | 0.00000 | 0.00000 | 0.00000 |
| Veillonellaceae | Dialister | 0.01984 | 0.02110 | 0.06983 | 0.00000 | 0.00000 | 0.00000 |
| Campylobacteraceae | Campylobacter | 0.01984 | 0.02275 | NA | 0.00000 | 0.00000 | 0.00000 |
| Tissierellaceae | Finegoldia | 0.04797 | 0.04647 | NA | 0.00000 | 0.00000 | 0.00001 |
| Porphyromonadaceae | Porphyromonas | 0.13031 | 0.10538 | NA | 0.00000 | 0.00000 | 0.00000 |
| Streptococcaceae | Streptococcus | 0.01984 | 0.02275 | NA | 0.00000 | 0.00000 | 0.00000 |
| Actinomycetaceae | Varibaculum | 0.55456 | 0.34057 | NA | 0.00001 | 0.28831 | 0.93412 |
| Prevotellaceae | Prevotella | 0.01984 | 0.02110 | NA | 0.00000 | 0.00000 | 0.00000 |
| Tissierellaceae | 1-68 | 0.84255 | 0.67380 | NA | 0.00006 | 0.00165 | 0.00171 |
| Lactobacillaceae | WAL 1855D | 0.09401 | 0.05409 | NA | 0.00000 | 0.00000 | 0.00000 |
| Tissierellaceae | Mobil uncus | 0.01984 | 0.02110 | NA | 0.00000 | 0.00000 | 0.00000 |
| Tissierellaceae | Atopobium | 0.01984 | 0.02522 | NA | 0.00000 | 0.00000 | 0.00000 |
| Bifidobacteriaceae | Gardnerella | 0.99925 | 0.79316 | NA | 0.00014 | 0.53031 | 0.59088 |
| Actinomycetaceae | Actinomyces | 0.01984 | 0.02110 | NA | 0.00000 | 0.00000 | 0.00000 |

Figure 3 shows the distribution of OTUs under *Lactobacillus*. Lactobacillus has five leaf nodes and the relative abundances of all the leaf node m = 1, 2, 3, 4 and 5 were higher in the pregnant group. *Lactobacillus* was observed to be more abundant in the pregnant group than in the healthy group and the absence of vaginal *Lactobacillus* species can increase the risk of preterm delivery [31]. Supplementary Figures 11–17 showed the OTU distributions of other associated genera. These results confirm that the genera identified using mTMAT may be associated with delivery. Therefore, mTMAT successfully detected associated genera.

**Figure 3.**
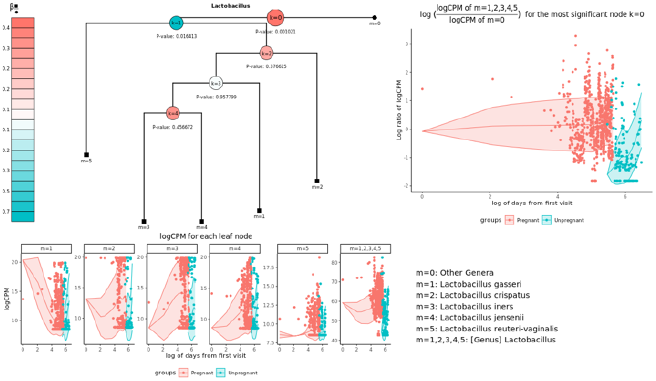
OTU distributions of significantly associated genus *Lactobacillus*. Relative proportions of OTUs belonging to *Lactobacillus* at different time points were plotted. Each OTU has its corresponding leaf node and leaf nodes in ▪ and ● indicate that they are in *L*_*k*_ and *R*_*k*_, respectively. 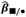 indicates the mean difference of *log(C*^*k*^_*ij*_*/D*^*k*^_*ij*_) between pregnant and unpregnant subjects after adjusting for covariates, and the red internal node indicates that OTUs in the left test leaf nodes are more abundant in pregnant subjects. The most significant node is enlarged.

## Discussion

The importance of microbiome-host interactions has been known for more than a century [32], and it has been shown that the occurrence of many human diseases is related to bacterial communities.

Microbiome data has phylogenetic structure and some unique properties, including high dimensionality, rarity and heterogeneity beyond composition. These properties cause multiple statistical problems when analyzing data across microbial composition and integrating multi-omics data such as large p and small n, dependencies, over-dispersion and zero inflation. The classical correlation and related methods throughout the microbial association study were applied in the actual study and used in the development of new methods. However, owing to those problems related to metagenomic analysis, traditional approaches are infrequently utilized for more complex models, such as longitudinal models including linear mixed models and generalized linear mixed models. Furthermore, those methods do not properly address compositional bias and may lead to erroneous results due to limitations in relative abundance data [33].

Here, we propose a new method for detecting OTUs associated with host diseases. mTMAT statistics are based on quasi-scores for internal nodes in a phylogenetic tree. It can take into account various correlation structures and provide robust estimation for mis-specified correlation structures while maintaining statistical validity for small sample size with the best statistical power. This property is achieved by using log CPM transformation and integrating abundances based on the phylogenetic tree. Correction of compositional bias is alleviated by taking ratios between two sets of integrated abundances.

These statistics are then combined to form a single statistic with the minimal p-value. By using such quasi-score statistics, mTMAT can identify differences among OTUs significantly associated with host disease status. Based on the nature of the proposed statistics, the statistical scores for internal nodes are independent, and the minimum p-value can be calculated directly. We compared the performance of mTMAT with those of GLMM-MiRKAT, LMM-arcsine, LMM-log, and FZINBMM under various simulation scenarios. According to our results, mTMAT correctly controlled the nominal type-1 error rate and was statistically the most powerful method for detecting associations with host diseases in our simulation studies. Additionally, methodologies using permutation-based p-values, such as GLMM-MiRKAT, are computationally very slow compared to mTMAT.

However, in spite of its usefulness, mTMAT has several limitations. First, mTMAT combines the statistics for each internal node, and multiple comparison occurs when the number of leaf nodes is large. Second, we evaluated the performance of the proposed methods with extensive simulations, but this result cannot be generalized. Third, the choice of database and OTU clustering methods can affect the statistical property of mTMAT. In our previous research, we showed that the effect of a mis-specified phylogenetic tree is generally not substantial. However, further investigation is still necessary, which we will undertake in our follow-up research.

mTMAT can help researchers easily perform fast and effective microbiome-wide association analysis, thereby comprehensively elucidating the interaction mechanism of the entire microbiota with the human body.

## References

Balvočiūtė, M. and Huson, D.H. SILVA, RDP, Greengenes, NCBI and OTT—how do these taxonomies compare? BMC genomics 2017;18(2):114.

Caporaso, J.G., et al. QIIME allows analysis of high-throughput community sequencing data. Nature methods 2010;7(5):335–336.

Chen, J., et al. Associating microbiome composition with environmental covariates using generalized UniFrac distances. Bioinformatics 2012;28(16):2106–2113.

Chun, J., et al. EzTaxon: a web-based tool for the identification of prokaryotes based on 16S ribosomal RNA gene sequences. International journal of systematic and evolutionary microbiology 2007;57(10):2259–2261.

de Vos, W.M. and de Vos, E.A. Role of the intestinal microbiome in health and disease: from correlation to causation. Nutrition reviews 2012;70(suppl_1):S45–S56.

Dethlefsen, L., McFall-Ngai, M. and Relman, D.A. An ecological and evolutionary perspective on human–microbe mutualism and disease. Nature 2007;449(7164):811.

Giloteaux, L., et al. Reduced diversity and altered composition of the gut microbiome in individuals with myalgic encephalomyelitis/chronic fatigue syndrome. Microbiome 2016;4(1):30.

Guo, S., et al. A Simple and Novel Fecal Biomarker for Colorectal Cancer: Ratio of Fusobacterium Nucleatum to Probiotics Populations, Based on Their Antagonistic Effect. Clinical chemistry 2018:clinchem. 2018.289728.

Janda, J.M. and Abbott, S.L. The genus Aeromonas: taxonomy, pathogenicity, and infection. Clinical microbiology reviews 2010;23(1):35–73.

Koh, H., Blaser, M.J. and Li, H. A powerful microbiome-based association test and a microbial taxa discovery framework for comprehensive association mapping. Microbiome 2017;5(1):45.

Law, C.W., et al. Voom: precision weights unlock linear model analysis tools for RNA-seq read counts. Genome biology 2014;15(2):R29.

Li, K., Bihan, M. and Methé, B.A. Analyses of the stability and core taxonomic memberships of the human microbiome. PloS one 2013;8(5):e63139.

Morgan, X.C., et al. Dysfunction of the intestinal microbiome in inflammatory bowel disease and treatment. Genome biology 2012;13(9):R79.

Park, S.-C. and Won, S. Evaluation of 16S rRNA Databases for Taxonomic Assignments Using Mock Community. Genomics & informatics 2018;16(4):e24.

Pielou, E.C. An introduction to mathematical ecology. An introduction to mathematical ecology. 1969.

Pruesse, E., Peplies, J. and Glöckner, F.O. SINA: accurate highthroughput multiple sequence alignment of ribosomal RNA genes. Bioinformatics 2012;28(14):1823–1829.

Quast, C., et al. The SILVA ribosomal RNA gene database project: improved data processing and web-based tools. Nucleic acids research 2012;41(D1):D590–D596.

Schloss, P.D., et al. Introducing mothur: open-source, platform-independent, community-supported software for describing and comparing microbial communities. Applied and environmental microbiology 2009;75(23):7537–7541.

Seelam, N.S., et al. Production, characterization and optimization of fermented tomato and carrot juices by using Lysinibacillus sphaericus isolate. Journal of Applied Biology & Biotechnology Vol 2017;5(04):066–075.

Wu, C., et al. An adaptive association test for microbiome data. Genome medicine 2016;8(1):56.

Zhang, X., et al. The oral and gut microbiomes are perturbed in rheumatoid arthritis and partly normalized after treatment. Nature medicine 2015;21(8):895–905.

Zhao, N., et al. Testing in microbiome-profiling studies with MiRKAT, the microbiome regression-based kernel association test. The American Journal of Human Genetics 2015;96(5):797–807.

