## Supplementary Text, Tables and Figures. for "Robust Phylogenetic Tree-based Microbiome Association Test using Repeatedly Measured Data for Composition Bias": mTMAT_Supp_May_2023.pdf

### Additional File 1: Supplementary Figures and Tables

**Supplementary Table 1. Type-1 error estimates of $mTMAT_M$ with genera from a longitudinal dataset. The values 1:1 and 1:3 were assumed for the ratio of cases and controls. The total sample size is denoted by N, and we considered N = 30, 50, and 100. All subjects were selected without replacement. Type-1 error estimates were calculated with 2,000 replicates at the significance levels 0.1, 0.05, 0.01, and 0.005. Compound symmetry (CS), first-order autoregressive (AR1), and unstructured (UN).**

| | Working Correlation matrix | Case : Control = 1 : 1 | | | Case : Control = 1 : 3 | | |
|---|---|---|---|---|---|---|---|
| | | N = 30 | N = 50 | N = 100 | N = 30 | N = 50 | N = 100 |
| α = 0.1 | Identity | 0.1061 | 0.1276 | 0.1621 | 0.1016 | 0.1181 | 0.1087 |
| | CS | 0.1034 | 0.1210 | 0.1518 | 0.1036 | 0.1159 | 0.1073 |
| | AR1 | 0.1058 | 0.1277 | 0.1602 | 0.1009 | 0.1157 | 0.1082 |
| | UN | 0.1007 | 0.1196 | 0.1443 | 0.1009 | 0.1154 | 0.1096 |
| α = 0.05 | Identity | 0.0493 | 0.0662 | 0.0940 | 0.0540 | 0.0617 | 0.0587 |
| | CS | 0.0487 | 0.0644 | 0.0859 | 0.0535 | 0.0579 | 0.0587 |
| | AR1 | 0.0490 | 0.0689 | 0.0939 | 0.0527 | 0.0590 | 0.0587 |
| | UN | 0.0479 | 0.0613 | 0.0809 | 0.0530 | 0.0576 | 0.0562 |
| α = 0.01 | Identity | 0.0085 | 0.0151 | 0.0266 | 0.0157 | 0.0166 | 0.0118 |
| | CS | 0.0084 | 0.0137 | 0.0222 | 0.0146 | 0.0153 | 0.0119 |
| | AR1 | 0.0081 | 0.0156 | 0.0259 | 0.0143 | 0.0150 | 0.0117 |
| | UN | 0.0084 | 0.0128 | 0.0199 | 0.0135 | 0.0151 | 0.0114 |
| α = 0.005 | Identity | 0.0040 | 0.0080 | 0.0156 | 0.0080 | 0.0092 | 0.0064 |
| | CS | 0.0041 | 0.0074 | 0.0130 | 0.0074 | 0.0082 | 0.0064 |
| | AR1 | 0.0043 | 0.0081 | 0.0150 | 0.0076 | 0.0084 | 0.0065 |
| | UN | 0.0037 | 0.0072 | 0.0112 | 0.0070 | 0.0082 | 0.0061 |

**Supplementary Table 2. Effect of numbers of leaf nodes on type-1 error estimates. Families were categorized into four different groups according to the number of leaf nodes, and for each taxon, type-1 error rates were estimated. Simulation data were generated with read counts from the dataset. We assumed the total sample size (N) was equal to 50. The value 1:3 was assumed for the ratio of cases and controls. Identity working correlation matrix and robust score statistics were used for mTMAT.**

| Method | Number of leaf nodes | Number of Family | Significance level | | | |
|---|---|---|---|---|---|---|
| | | | α = 0.1 | α = 0.05 | α = 0.01 | α = 0.005 |
| $mTMAT_{IM}$ | 1 | 22 | 0.1050 | 0.0508 | 0.0102 | 0.0047 |
| | 2-5 | 12 | 0.0917 | 0.0411 | 0.0042 | 0.0014 |
| | 6-15 | 5 | 0.0733 | 0.0211 | 0.0000 | 0.0000 |
| | >15 | 2 | 0.0700 | 0.0267 | 0.0017 | 0.0017 |
| $mTMAT_{M}$ | 1 | 22 | 0.1147 | 0.0641 | 0.0182 | 0.0112 |
| | 2-5 | 12 | 0.1014 | 0.0442 | 0.0078 | 0.0039 |
| | 6-15 | 5 | 0.0833 | 0.0433 | 0.0044 | 0.0022 |
| | >15 | 2 | 0.0700 | 0.0267 | 0.0033 | 0.0017 |
| GLMM-MiRKAT | 1 | 22 | NA | NA | NA | NA |
| | 2-5 | 12 | 0.1508 | 0.0872 | 0.0222 | 0.0125 |
| | 6-15 | 5 | 0.2322 | 0.1444 | 0.0433 | 0.0244 |
| | >15 | 2 | 0.2633 | 0.1933 | 0.0717 | 0.0417 |
| FZINBMM | 1 | 22 | 0.5683 | 0.5092 | 0.4035 | 0.3717 |
| | 2-5 | 12 | 0.2806 | 0.2147 | 0.1228 | 0.0997 |
| | 6-15 | 5 | 0.1133 | 0.0633 | 0.0167 | 0.0111 |
| | >15 | 2 | 0.1000 | 0.0333 | 0.0033 | 0.0000 |
| LMM-arcsine | 1 | 22 | 0.0549 | 0.0315 | 0.0067 | 0.0042 |
| | 2-5 | 12 | 0.0947 | 0.0450 | 0.0101 | 0.0054 |
| | 6-15 | 5 | 0.1211 | 0.0589 | 0.0133 | 0.0034 |
| | >15 | 2 | 0.1633 | 0.1017 | 0.0267 | 0.0133 |
| LMM-log | 1 | 22 | 0.0920 | 0.0414 | 0.0118 | 0.0059 |
| | 2-5 | 12 | 0.991 | 0.0509 | 0.0095 | 0.0048 |
| | 6-15 | 5 | 0.0971 | 0.0426 | 0.0123 | 0.0045 |
| | >15 | 2 | 0.1117 | 0.0687 | 0.0067 | 0.0050 |

**Supplementary Table 3. Effect of sparsity on type-1 error estimates. For each genus, we calculated its sparsity as the proportion of subjects with no abundance. Genera were sorted by their sparsity and categorized into three different groups, and for each taxon, type-1 error rates were estimated. Simulation data were generated by using read counts from the dataset. We assumed the total sample size (N) was equal to 50. The value 1:3 was assumed for the ratio of cases and controls. Identity working correlation matrix and robust score statistics were used for mTMAT.**

| Method | Mean sparsity level of genera | Mean number of leaf nodes | Number of genus | Significance level | | | |
|---|---|---|---|---|---|---|---|
| | | | | α = 0.1 | α = 0.05 | α = 0.01 | α = 0.005 |
| $mTMAT_{IM}$ | <=20% | 1 | 3 | 0.0989 | 0.0522 | 0.0056 | 0.0022 |
| | 20-50% | 2.25 | 12 | 0.0883 | 0.0417 | 0.0061 | 0.0019 |
| | >50% | 1.66 | 58 | 0.1032 | 0.0496 | 0.0090 | 0.0039 |
| $mTMAT_{M}$ | <=20% | 1 | 3 | 0.1200 | 0.0600 | 0.0111 | 0.0067 |
| | 20-50% | 2.25 | 12 | 0.0989 | 0.0447 | 0.0072 | 0.0042 |
| | >50% | 1.66 | 58 | 0.1180 | 0.0620 | 0.0152 | 0.0093 |
| GLMM-MiRKAT | <=20% | 1 | 3 | NA | NA | NA | NA |
| | 20-50% | 2.25 | 12 | 0.1225 | 0.0625 | 0.0100 | 0.0025 |
| | >50% | 1.66 | 58 | 0.1526 | 0.1047 | 0.0611 | 0.0584 |
| FZINBMM | <=20% | 1 | 3 | 0.2067 | 0.1333 | 0.0500 | 0.0367 |
| | 20-50% | 2.25 | 12 | 0.2600 | 0.1900 | 0.1083 | 0.0858 |
| | >50% | 1.66 | 58 | 0.4961 | 0.4351 | 0.3297 | 0.2964 |
| LMM-arcsine | <=20% | 1 | 3 | 0.1013 | 0.0491 | 0.0045 | 0.0067 |
| | 20-50% | 2.25 | 12 | 0.0973 | 0.0482 | 0.0132 | 0.0047 |
| | >50% | 1.66 | 58 | 0.0828 | 0.0381 | 0.0084 | 0.0051 |
| LMM-log | <=20% | 1 | 3 | 0.0956 | 0.0501 | 0.0043 | 0.0032 |
| | 20-50% | 2.25 | 12 | 0.0962 | 0.0434 | 0.0078 | 0.0041 |
| | >50% | 1.66 | 58 | 0.0972 | 0.0499 | 0.0112 | 0.0063 |

**Supplementary Table 4. Effect of assumed correlation structure on type-1 error estimates. For each taxon, type-1 error rates were estimated. Simulation data were generated by using read counts from the simulation dataset with microbiomeDASim package. We assumed the total sample size (N) was equal to 50. Identity working correlation matrix and robust score statistics were used for mTMAT$_{IM}$. The number of time points was set to 6.**

| Assumed Correlation Structure | Assumed rho | Working Correlation Structure | Significance level | | | |
|---|---|---|---|---|---|---|
| | | | α = 0.1 | α = 0.05 | α = 0.01 | α = 0.005 |
| Identity | 0 | Identity | 0.1045 | 0.0473 | 0.0078 | 0.0033 |
| | | CS | 0.1045 | 0.0503 | 0.0103 | 0.0035 |
| | | AR1 | 0.1035 | 0.0515 | 0.0100 | 0.0040 |
| | | Unstructured | 0.1043 | 0.0515 | 0.0100 | 0.0040 |
| CS | 0.2 | Identity | 0.1020 | 0.0520 | 0.0095 | 0.0048 |
| | | CS | 0.0998 | 0.0493 | 0.0085 | 0.0030 |
| | | AR1 | 0.0983 | 0.0488 | 0.0080 | 0.0030 |
| | | Unstructured | 0.0995 | 0.0505 | 0.0070 | 0.0033 |
| | 0.5 | Identity | 0.1028 | 0.0455 | 0.0090 | 0.0053 |
| | | CS | 0.0953 | 0.0463 | 0.0083 | 0.0043 |
| | | AR1 | 0.0970 | 0.0445 | 0.0078 | 0.0048 |
| | | Unstructured | 0.0955 | 0.0453 | 0.0080 | 0.0045 |
| | 0.8 | Identity | 0.0988 | 0.0483 | 0.0095 | 0.0050 |
| | | CS | 0.0980 | 0.0488 | 0.0073 | 0.0033 |
| | | AR1 | 0.0973 | 0.0493 | 0.0073 | 0.0033 |
| | | Unstructured | 0.0958 | 0.0478 | 0.0075 | 0.0033 |
| AR1 | 0.2 | Identity | 0.1013 | 0.0523 | 0.0123 | 0.0068 |
| | | CS | 0.0983 | 0.0465 | 0.0073 | 0.0035 |
| | | AR1 | 0.0975 | 0.0463 | 0.0080 | 0.0035 |
| | | Unstructured | 0.0958 | 0.0475 | 0.0070 | 0.0033 |
| | 0.5 | Identity | 0.1000 | 0.0510 | 0.0100 | 0.0045 |
| | | CS | 0.0993 | 0.0505 | 0.0083 | 0.0035 |
| | | AR1 | 0.0993 | 0.0510 | 0.0090 | 0.0033 |
| | | Unstructured | 0.1005 | 0.0480 | 0.0078 | 0.0033 |
| | 0.8 | Identity | 0.1015 | 0.0468 | 0.0060 | 0.0015 |
| | | CS | 0.0913 | 0.0405 | 0.0083 | 0.0038 |
| | | AR1 | 0.0913 | 0.0395 | 0.0080 | 0.0035 |
| | | Unstructured | 0.0923 | 0.0398 | 0.0065 | 0.0038 |

**Supplementary Table 5. Type-1 error and the power estimates of disease and time effect when the interaction between time and disease group does not exist. Simulation data were generated by using microbiomeDASim package. We assumed the total sample size (N) was equal to 50. Identity working correlation matrix and robust score statistic are used for $mTMAT_{IM}$ and $mTMAT_{M}$. The number of time points was set to 6. The significance level was set to 0.05. The working correlation was set to CS and the rho was set to 0.1. The power estimates for the time effect of GLMM-MiRKAT were set to be NA because the time effect for the model FZINBMM was excluded, as FZINBMM had a singularity problem for more than 90% of the simulated datasets.**

| Method | Assumptions | Null hypothesis | |
|---|---|---|---|
| | | $\beta = 0$ | $\beta_{time} = 0$ |
| $mTMAT_{IM}$ | $\beta = 0.1, \beta_{time} = 0$ | 0.05 | 0.05 |
| $mTMAT_{IM}$ | $\beta = 0.5, \beta_{time} = 0$ | 0.08 | 0.04 |
| $mTMAT_{IM}$ | $\beta = 1, \beta_{time} = 0$ | 0.48 | 0.05 |
| $mTMAT_{IM}$ | $\beta = 0, \beta_{time} = 0.1$ | 0.04 | 0.12 |
| $mTMAT_{IM}$ | $\beta = 0, \beta_{time} = 0.5$ | 0.03 | 0.50 |
| $mTMAT_{IM}$ | $\beta = 0, \beta_{time} = 1$ | 0.03 | 0.99 |
| $mTMAT_{M}$ | $\beta = 0.1, \beta_{time} = 0$ | 0.04 | 0.04 |
| $mTMAT_{M}$ | $\beta = 0.5, \beta_{time} = 0$ | 0.07 | 0.04 |
| $mTMAT_{M}$ | $\beta = 1, \beta_{time} = 0$ | 0.46 | 0.05 |
| $mTMAT_{M}$ | $\beta = 0, \beta_{time} = 0.1$ | 0.05 | 0.11 |
| $mTMAT_{M}$ | $\beta = 0, \beta_{time} = 0.5$ | 0.04 | 0.47 |
| $mTMAT_{M}$ | $\beta = 0, \beta_{time} = 1$ | 0.03 | 0.97 |
| GLMM-MiRKAT | $\beta = 0.1, \beta_{time} = 0$ | 0.80 | NA |
| GLMM-MiRKAT | $\beta = 0.5, \beta_{time} = 0$ | 0.78 | NA |
| GLMM-MiRKAT | $\beta = 1, \beta_{time} = 0$ | 0.77 | NA |
| GLMM-MiRKAT | $\beta = 0, \beta_{time} = 0.1$ | 0.81 | NA |
| GLMM-MiRKAT | $\beta = 0, \beta_{time} = 0.5$ | 0.78 | NA |
| GLMM-MiRKAT | $\beta = 0, \beta_{time} = 1$ | 0.77 | NA |
| LMM-arcsine | $\beta = 0.1, \beta_{time} = 0$ | 0.06 | 0.08 |
| LMM-arcsine | $\beta = 0.5, \beta_{time} = 0$ | 0.24 | 0.08 |
| LMM-arcsine | $\beta = 1, \beta_{time} = 0$ | 0.69 | 0.07 |
| LMM-arcsine | $\beta = 0, \beta_{time} = 0.1$ | 0.03 | 0.23 |
| LMM-arcsine | $\beta = 0, \beta_{time} = 0.5$ | 0.04 | 0.73 |
| LMM-arcsine | $\beta = 0, \beta_{time} = 1$ | 0.03 | 1 |
| LMM-log | $\beta = 0.1, \beta_{time} = 0$ | 0.05 | 0.09 |
| LMM-log | $\beta = 0.5, \beta_{time} = 0$ | 0.22 | 0.08 |
| LMM-log | $\beta = 1, \beta_{time} = 0$ | 0.69 | 0.09 |
| LMM-log | $\beta = 0, \beta_{time} = 0.1$ | 0.04 | 0.22 |
| LMM-log | $\beta = 0, \beta_{time} = 0.5$ | 0.03 | 0.72 |
| LMM-log | $\beta = 0, \beta_{time} = 1$ | 0.03 | 0.99 |

**Supplementary Table 6. Type-1 error and the power estimates of disease effect, time effect and interaction effect when the interaction between time and disease group exists. Simulation data were generated by using read counts from the simulation dataset with microbiomeDASim package. We assumed the total sample size (N) was equal to 50. Identity working correlation matrix and robust score statistics were used for $mTMAT_{IM}$ and $mTMAT_{M}$. The number of time points was set to 6. The significance level was set to 0.05. The working correlation was set to CS and the rho was set to 0.1. GLMM-MiRKAT and FZINBMM were excluded because GLMM-MiRKAT was not applicable to add time or interaction effects for the model and FZINBMM had a singularity problem for more than 90% of the simulated datasets.**

| Method | Assumptions | Null hypothesis | | |
|---|---|---|---|---|
| | | $\beta = 0$ | $\beta_{time} = 0$ | $\beta_{inter} = 0$ |
| $mTMAT_{IM}$ | $\beta = 0.1, \beta_{time} = 0.1, \beta_{inter} = 0$ | 0.05 | 0.12 | 0.05 |
| $mTMAT_{IM}$ | $\beta = 0.1, \beta_{time} = 0.1, \beta_{inter} = 0.1$ | 0.03 | 0.04 | 0.04 |
| $mTMAT_{IM}$ | $\beta = 0.1, \beta_{time} = 0.1, \beta_{inter} = 0.5$ | 0.04 | 0.05 | 0.13 |
| $mTMAT_{IM}$ | $\beta = 0.1, \beta_{time} = 0.1, \beta_{inter} = 1$ | 0.03 | 0.05 | 0.51 |
| $mTMAT_{M}$ | $\beta = 0.1, \beta_{time} = 0.1, \beta_{inter} = 0$ | 0.04 | 0.11 | 0.04 |
| $mTMAT_{M}$ | $\beta = 0.1, \beta_{time} = 0.1, \beta_{inter} = 0.1$ | 0.04 | 0.04 | 0.04 |
| $mTMAT_{M}$ | $\beta = 0.1, \beta_{time} = 0.1, \beta_{inter} = 0.5$ | 0.05 | 0.04 | 0.12 |
| $mTMAT_{M}$ | $\beta = 0.1, \beta_{time} = 0.1, \beta_{inter} = 1$ | 0.03 | 0.05 | 0.49 |
| LMM-arcsine | $\beta = 0.1, \beta_{time} = 0.1, \beta_{inter} = 0$ | 0.03 | 0.23 | 0.04 |
| LMM-arcsine | $\beta = 0.1, \beta_{time} = 0.1, \beta_{inter} = 0.1$ | 0.04 | 0.22 | 0.09 |
| LMM-arcsine | $\beta = 0.1, \beta_{time} = 0.1, \beta_{inter} = 0.5$ | 0.04 | 0.24 | 0.25 |
| LMM-arcsine | $\beta = 0.1, \beta_{time} = 0.1, \beta_{inter} = 1$ | 0.04 | 0.22 | 0.74 |
| LMM-log | $\beta = 0.1, \beta_{time} = 0.1, \beta_{inter} = 0$ | 0.03 | 0.21 | 0.05 |
| LMM-log | $\beta = 0.1, \beta_{time} = 0.1, \beta_{inter} = 0.1$ | 0.03 | 0.20 | 0.12 |
| LMM-log | $\beta = 0.1, \beta_{time} = 0.1, \beta_{inter} = 0.5$ | 0.04 | 0.22 | 0.26 |
| LMM-log | $\beta = 0.1, \beta_{time} = 0.1, \beta_{inter} = 1$ | 0.03 | 0.21 | 0.82 |

**A. Phase 1**

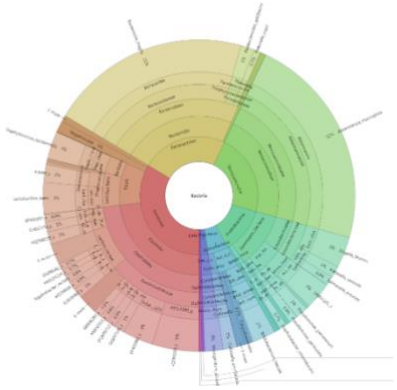

**B. Phase 2**

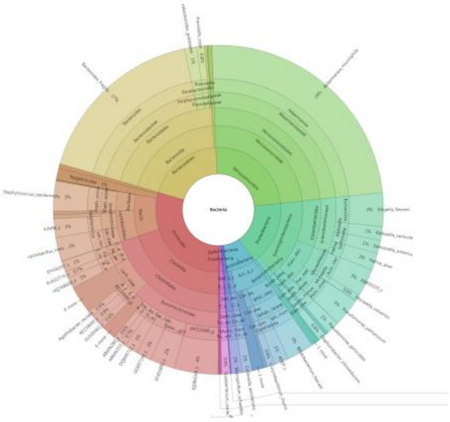

**C. Phase 3**

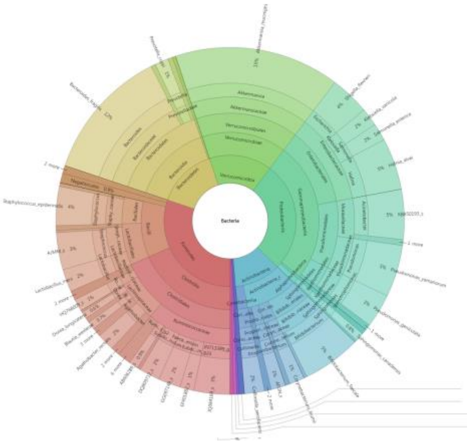

**Supplementary Figure 1. Taxonomic composition. Krona plots for phases 1, 2, and 3 showing the mean relative abundances of bacterial taxa at different taxonomic levels.**

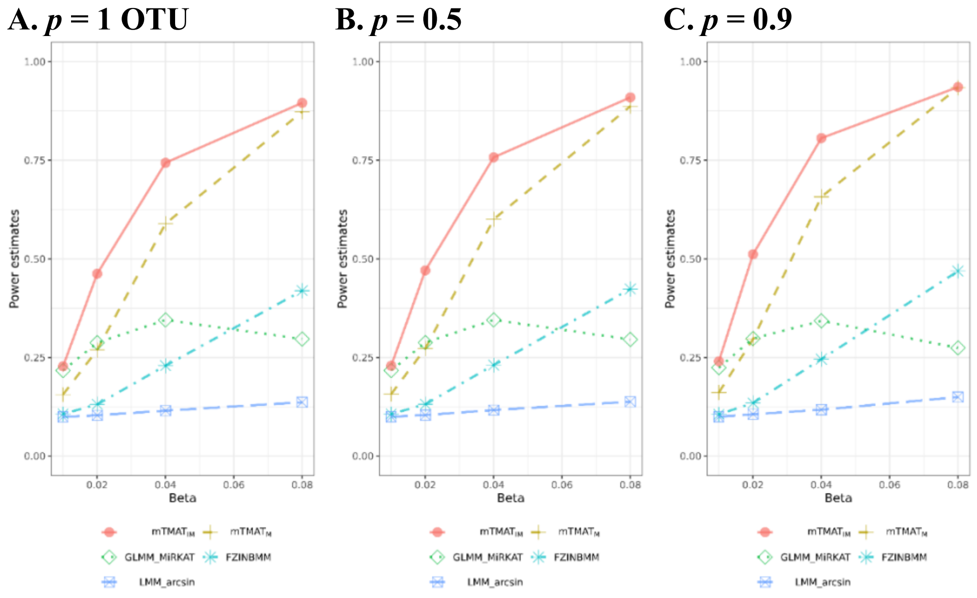


**Supplementary Figure 2. Power estimates for genera consisting of one or more operational taxonomic units (OTUs).** Power estimates at the significance level of 0.05 were calculated with 500 replicates. We generated simulation data based on read counts from datasets, and results from GLMM-MIRKAT were excluded because they cannot be applied to genera consisting of a single OTU. The significance levels for each method were adjusted to that of type-1 error rates based on the statistics from the simulation under the null hypothesis. We assumed the total sample size (N) to be equal to 50 and the ratio of cases and controls was set to 1:3 at a missing rate of 10%. The identity working correlation matrix and robust score statistics were used for mTMAT.

**A.** $p$ **= 1 OTU** **B.** $p$ **= 0.5** **C.** $p$ **= 0.9**

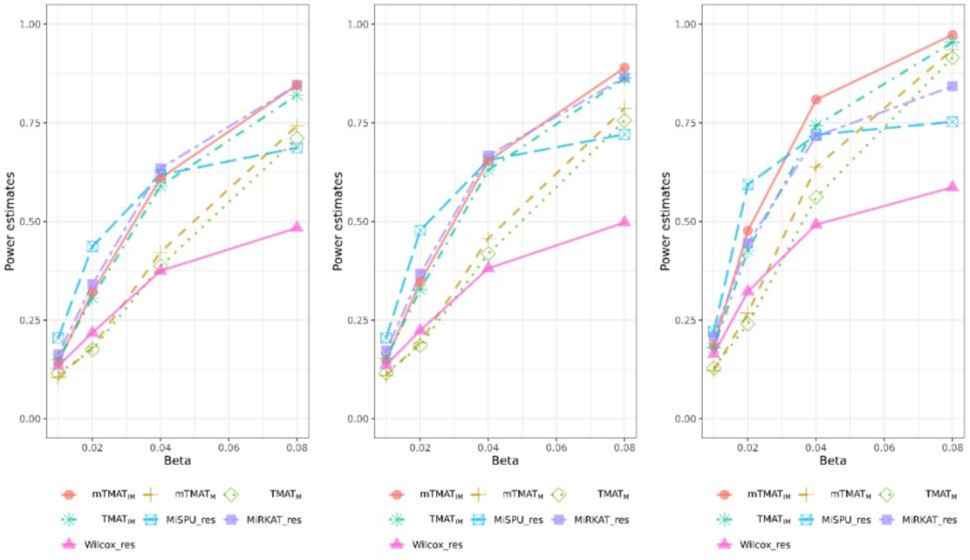


**Supplementary Figure 3. Comparison of power estimates with the methods for cross-sectionally observed data.** Power estimates at the significance level of 0.05 were calculated with 500 replicates. We assumed the total sample size (N) to be equal to 50 and the ratio of cases and controls was set to 1:3 at a missing rate 10%. Identity working correlation matrix and robust score statistics were used for mTMAT.

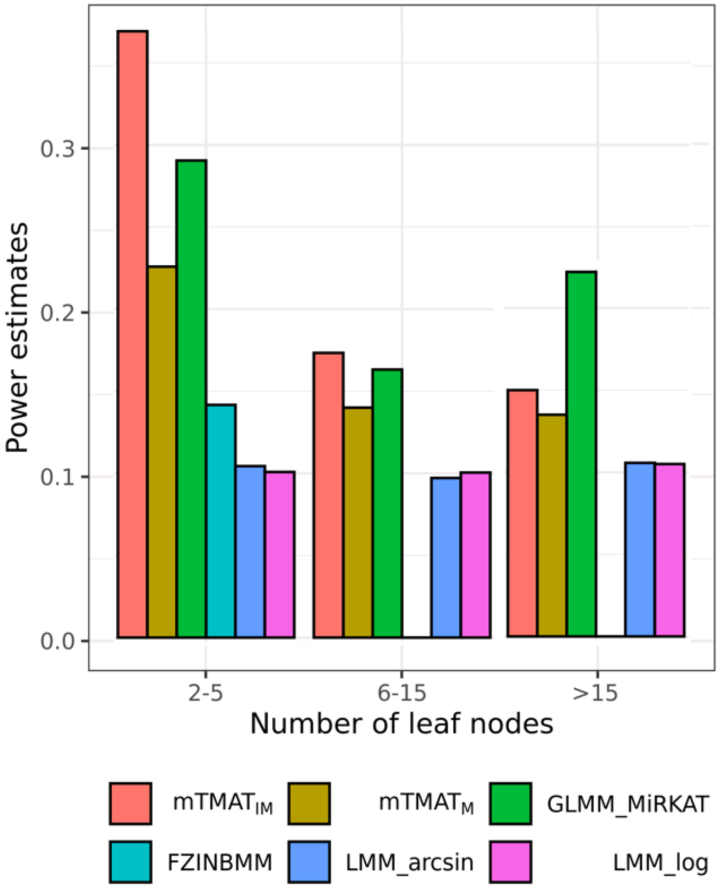


**Supplementary Figure 4. Effect of numbers of leaf nodes on power estimates.** Families were categorized into four different groups according to the number of leaf nodes, and for each taxon, power estimates at the 0.05 significance level were calculated with 500 replicates. We generated simulation data based on read counts from datasets, and the results were combined. We considered families with more than one OTU. We assumed the total sample size ($N$) = 50 at a missing rate of 10%, p = 50%, and $\boldsymbol{\beta}$ = 0.02; the ratio of cases and controls was set to 1:3 at a missing rate of 10%.

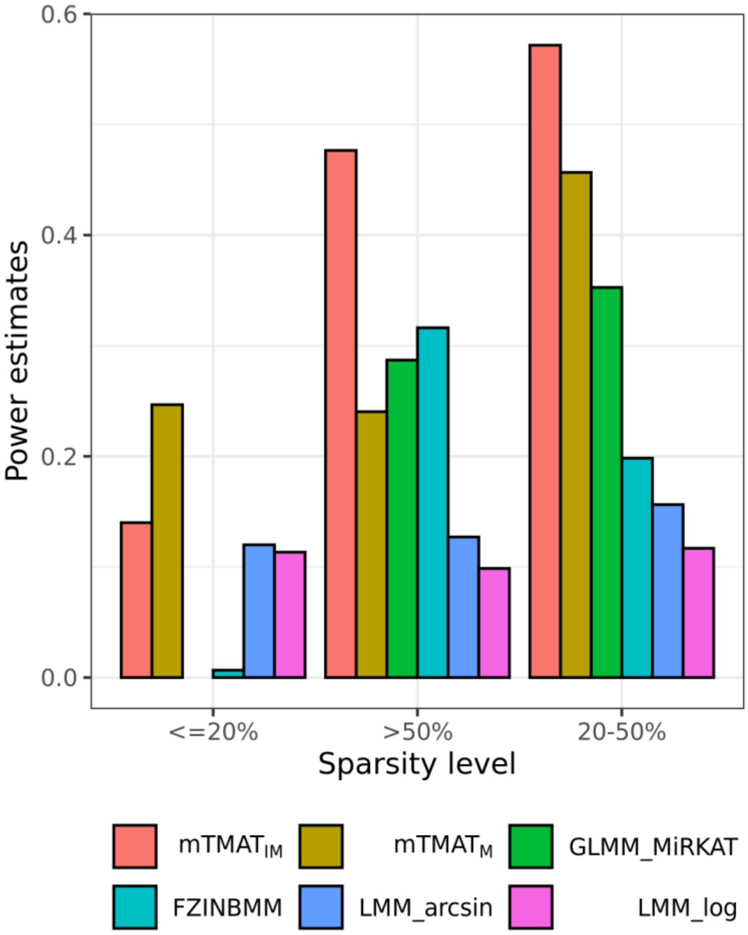


**Supplementary Figure 5. Effect of sparsity on power estimates.** We calculated the sparsity of each genus as the proportion of subjects with no reads (abundance of 0). Genera were sorted according to their sparsity and categorized into three different groups, and for each taxon, power estimates at the 0.05 significance level were calculated with 500 replicates. We generated simulation data based on read counts from the dataset and considered genera with more than one OTU. We assumed the total sample size ($N$) = 50 at a missing rate of 10%, p = 50%, and $\boldsymbol{\beta}$ = 0.02; the ratio of cases and controls was set as 1:3 at a missing rate of 10%.

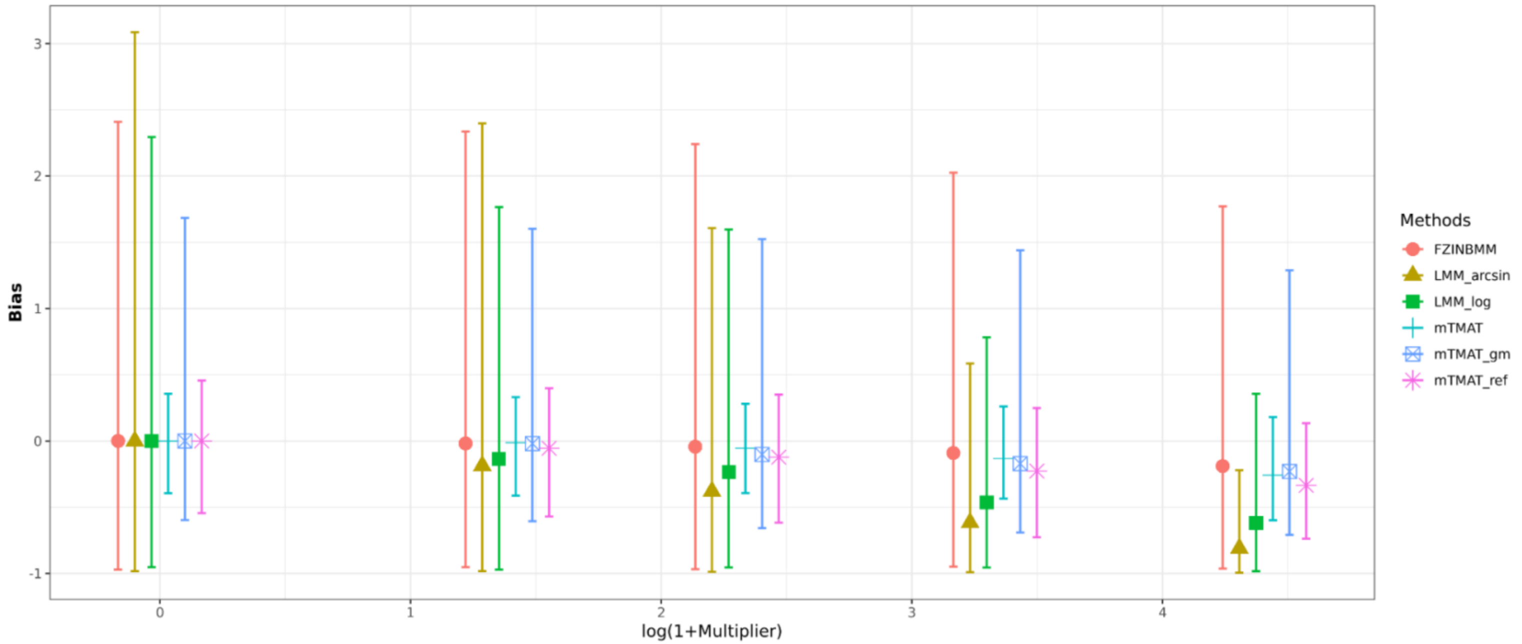


**Supplementary Figure 6. Effect of compositional bias for the methods.** Bias is calculated using the difference between beta and estimated. The bar represents the inter-quantile range of the bias. We performed 200 simulations assuming the total sample size (*N*) = 50, p = 50%, and $\beta$ = 0.15; the ratio of cases and controls was set to 1:3. $\mathrm{mTMAT_{gm}}$ and $\mathrm{mTMAT_{ref}}$ are two variations of mTMAT. $\mathrm{mTMAT_{gm}}$ uses the geometric mean of $E_{ij}^{t}$ for all taxonomies for the value of $G_{ij}$ while $\mathrm{mTMAT_{ref}}$ refers to a variation that uses $E_{ij}^{t}$ of a reference taxon for the value of $G_{ij}$. The most abundant taxon is chosen as the reference taxon.

**A. Baseline**

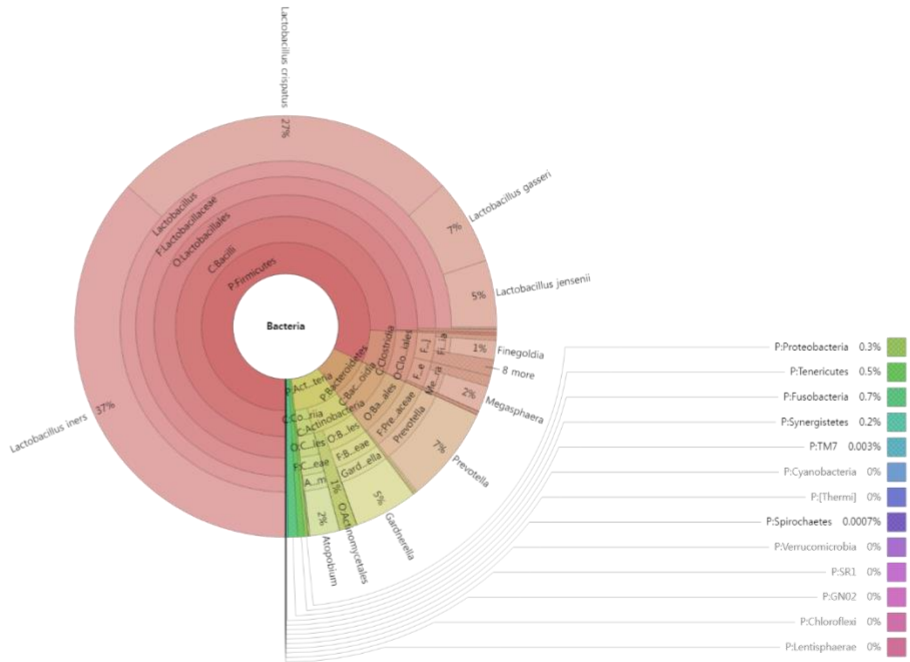


**B. 0-200 days from baseline**

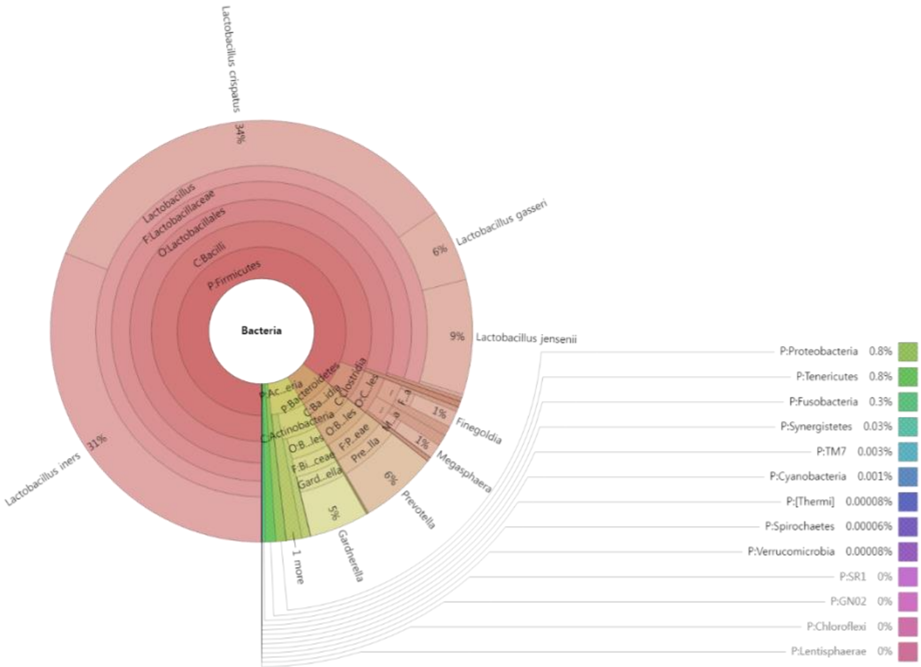


**Supplementary Figure 7. Change in microbial composition.** Krona plots showing the mean relative abundance of bacterial taxa with different time ranges at the species level. *(Continued)*

**C. 200-300 days from baseline**

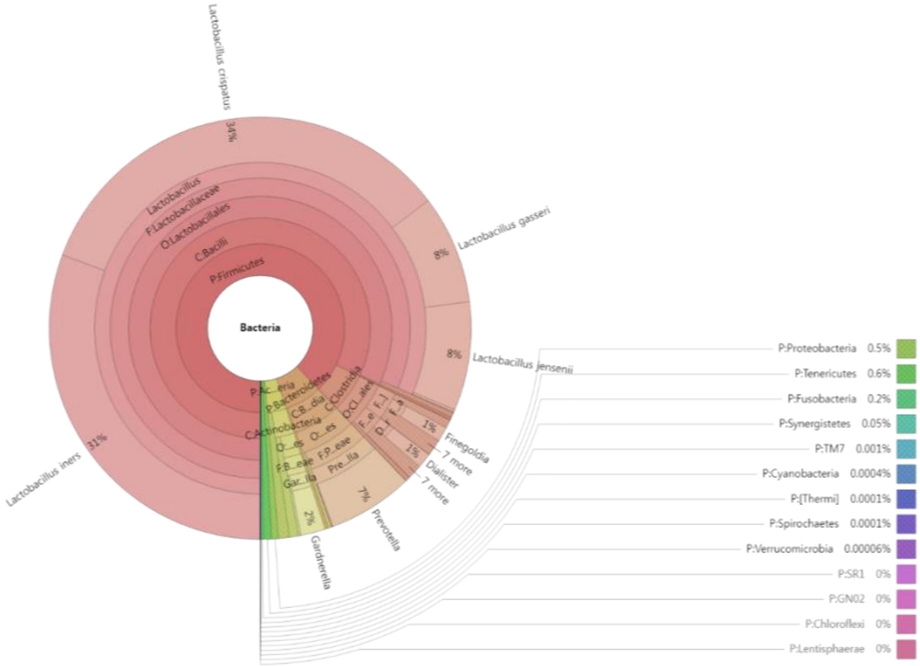


**D. More than 300 days from baseline**

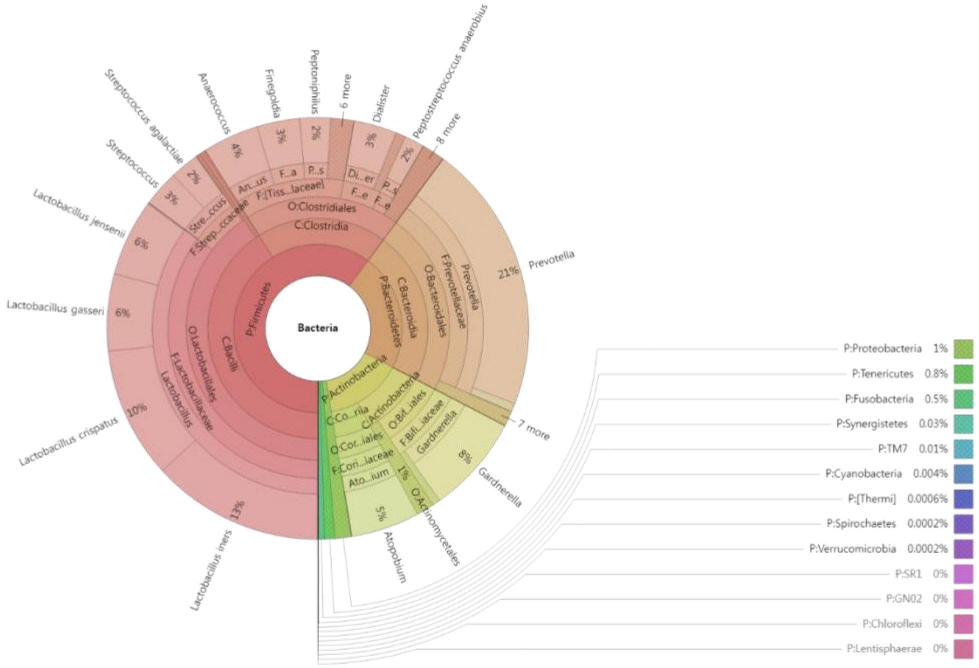


**Supplementary Figure 7. Continued**

**A. Pregnant**

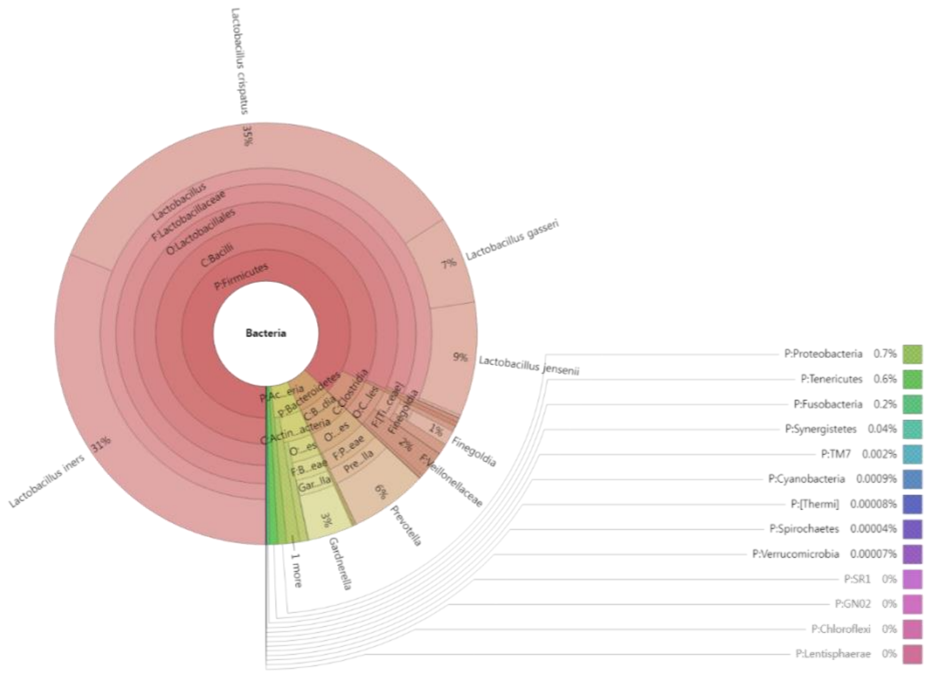


**B. Unpregnant**

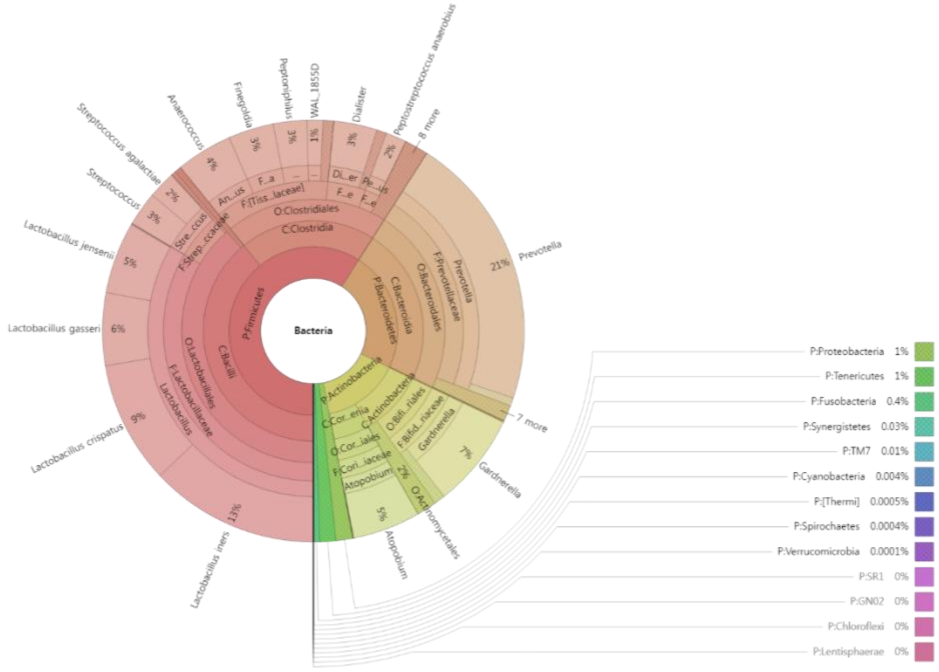


**Supplementary Figure 8. Microbial composition by pregnant groups.** Krona plots showing the mean relative abundance of bacterial taxa with different pregnant groups at the species level.

**A. Entire visit** **B. Baseline**

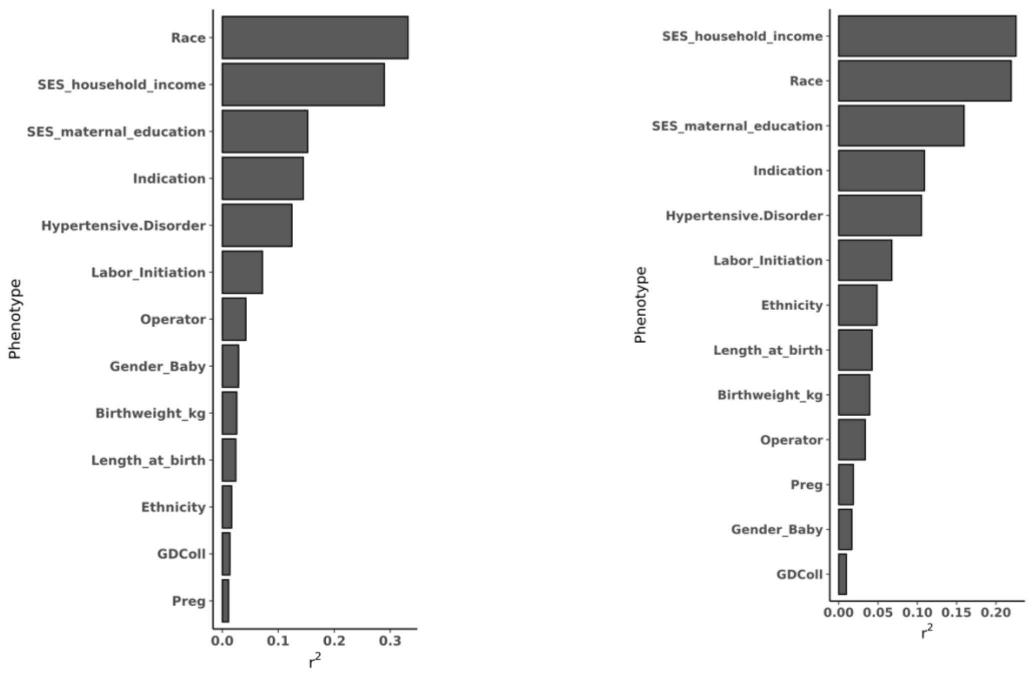


**Supplementary Figure 9. Relative importance of variables.** Relative proportions of variance attributable to each variable were calculated with PERMANOVA. Pldist and bray-curtis distance was used for the calculation of beta diversity.

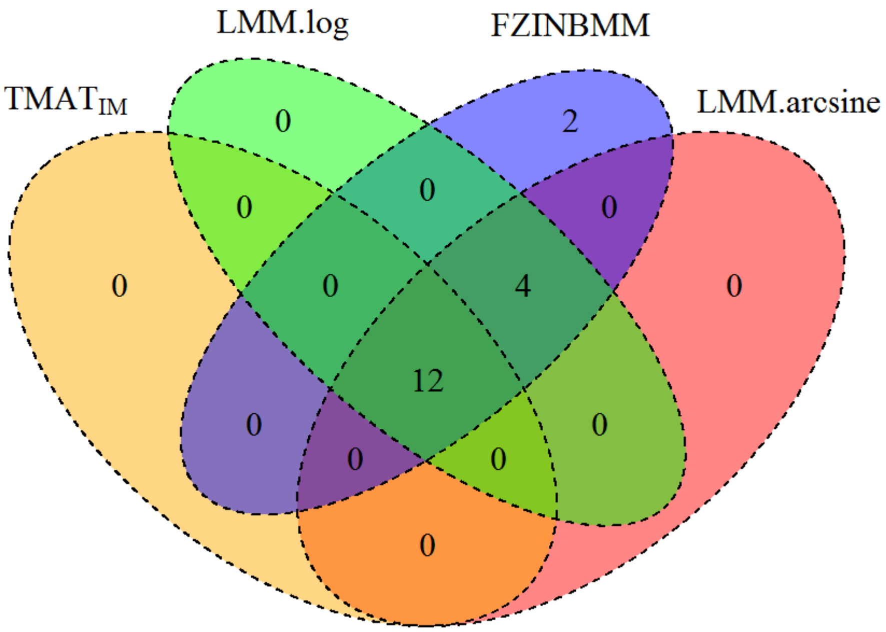


**Supplementary Figure 10. Comparison of significantly associated genera between different statistical methods.** Number of significantly associated genera at the FDR-adjusted 0.05 significance level are compared between different methods.

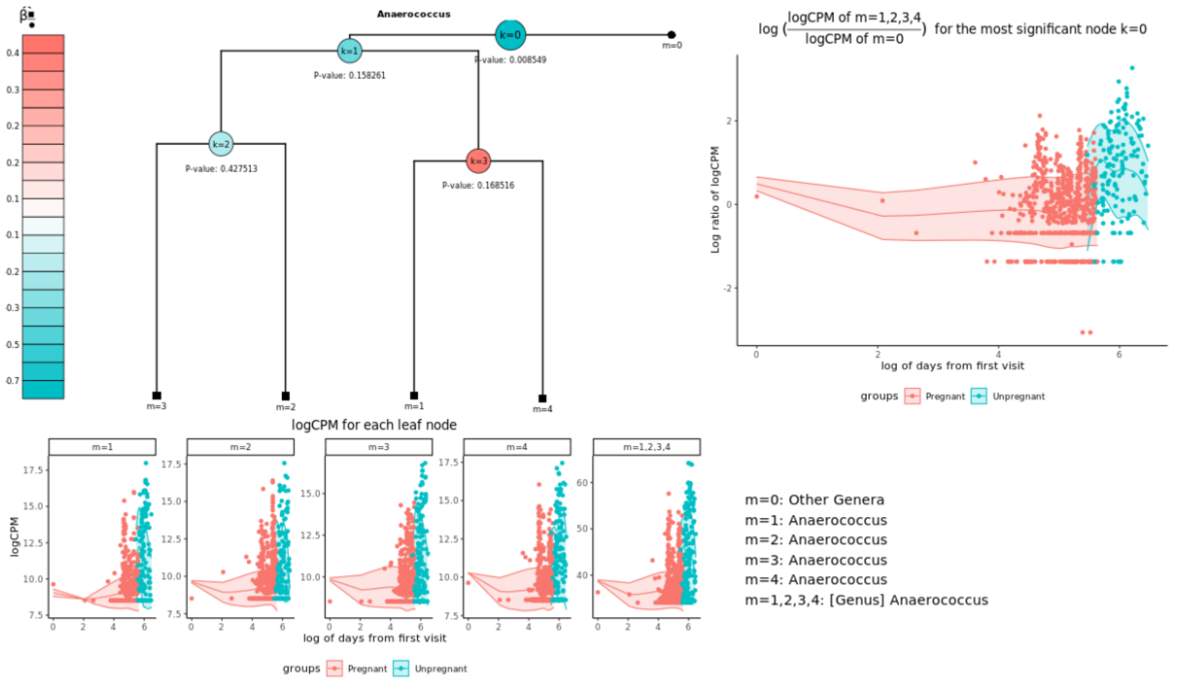


**Supplementary Figure 11. OTU distributions of significantly associated genus *Anaerococcus*.** Relative proportions of OTUs belonging to *Anaerococcus* at different time points were plotted. Each OTU has its corresponding leaf node and leaf nodes in ■ and ● indicate that they are in $\boldsymbol{L_k}$ and $\boldsymbol{R_k}$, respectively. $\hat{\boldsymbol{\beta}}_{\frac{\blacksquare}{\bullet}}$ indicates the mean difference of $log(C^k_{ij}/D^k_{ij})$ between pregnant and unpregnant subjects after adjusting for covariates, and the red internal node indicates that OTUs in the left test leaf nodes are more abundant in pregnant subjects. The most significant node is enlarged.

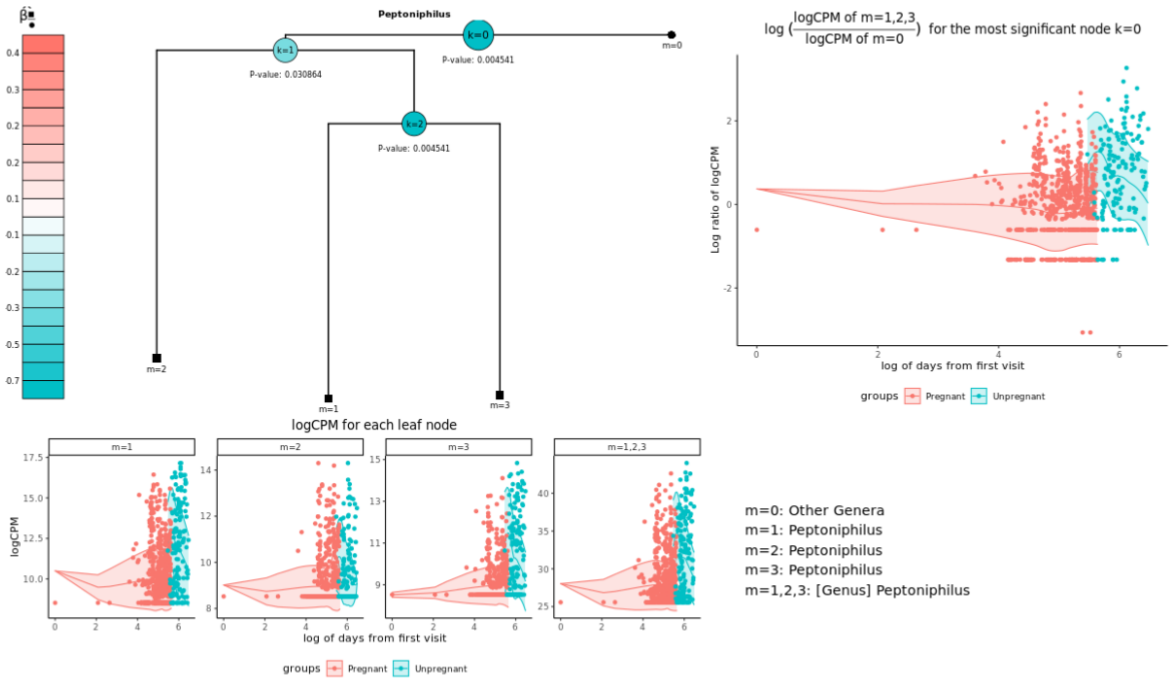


**Supplementary Figure 12. OTU distributions of significantly associated genus *Peptoniphilus*.** Relative proportions of OTUs belonging to *Peptoniphilus* at different time points were plotted. Each OTU has its corresponding leaf node and leaf nodes in ■ and ● indicate that they are in $\boldsymbol{L_k}$ and $\boldsymbol{R_k}$, respectively. $\hat{\boldsymbol{\beta}}_{\frac{\blacksquare}{\bullet}}$ indicates the mean difference of $log(C^k_{ij}/D^k_{ij})$ between pregnant and unpregnant subjects after adjusting for covariates, and the red internal node indicates that OTUs in the left test leaf nodes are more abundant in pregnant subjects. The most significant node is enlarged.

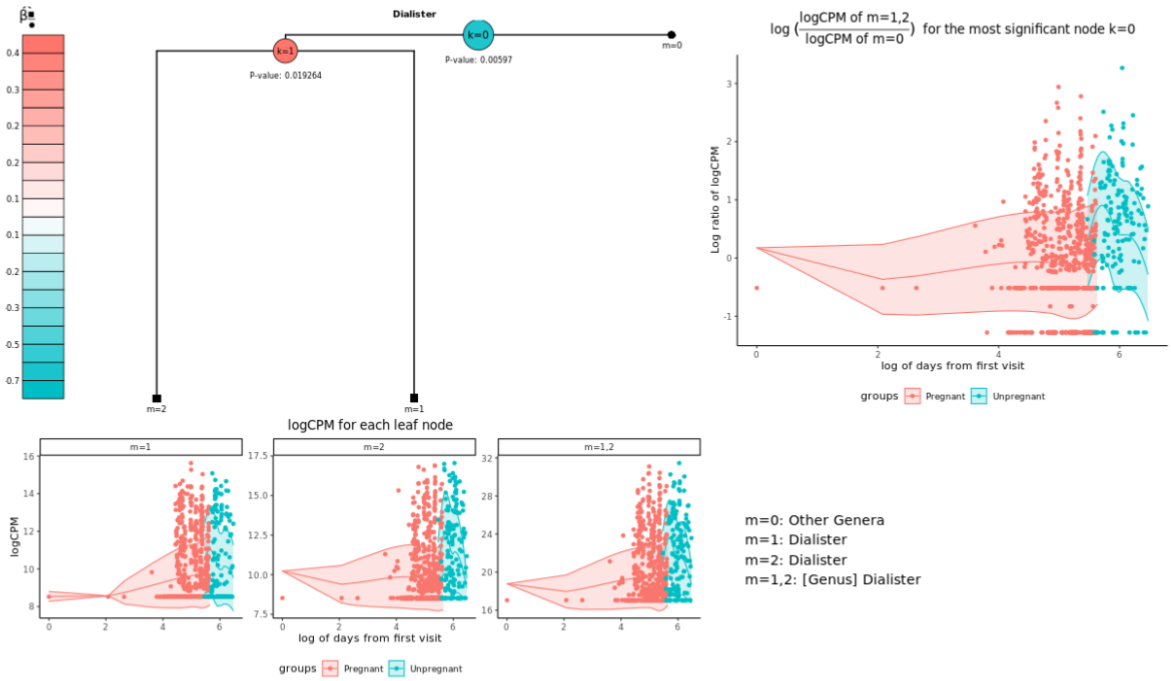


**Supplementary Figure 13. OTU distributions of significantly associated genus *Dialister*.** Relative proportions of OTUs belonging to *Dialister* at different time points were plotted. Each OTU has its corresponding leaf node and leaf nodes in ■ and ● indicate that they are in $\boldsymbol{L_k}$ and $\boldsymbol{R_k}$, respectively. $\widehat{\boldsymbol{\beta}}_{\frac{\blacksquare}{\bullet}}$ indicates the mean difference of $log(C^{k}_{ij}/D^{k}_{ij})$ between pregnant and unpregnant subjects after adjusting for covariates, and the red internal node indicates that OTUs in the left test leaf nodes are more abundant in pregnant subjects. The most significant node is enlarged.

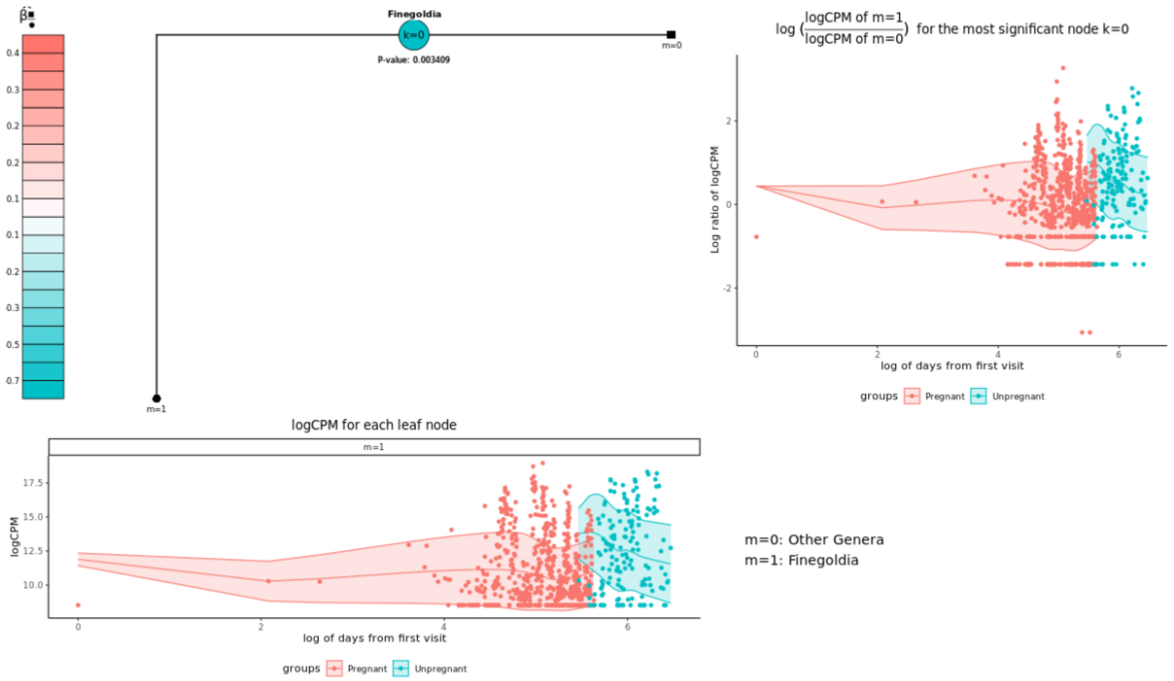


**Supplementary Figure 14. OTU distributions of significantly associated genus *Finegoldia*.** Relative proportions of OTUs belonging to *Finegoldia* at different time points were plotted. Each OTU has its corresponding leaf node and leaf nodes in ■ and ● indicate that they are in $\boldsymbol{L_k}$ and $\boldsymbol{R_k}$, respectively. $\widehat{\boldsymbol{\beta}}_{\frac{\blacksquare}{\bullet}}$ indicates the mean difference of $log(C^k_{ij}/D^k_{ij})$ between pregnant and unpregnant subjects after adjusting for covariates, and the red internal node indicates that OTUs in the left test leaf nodes are more abundant in pregnant subjects. The most significant node is enlarged.

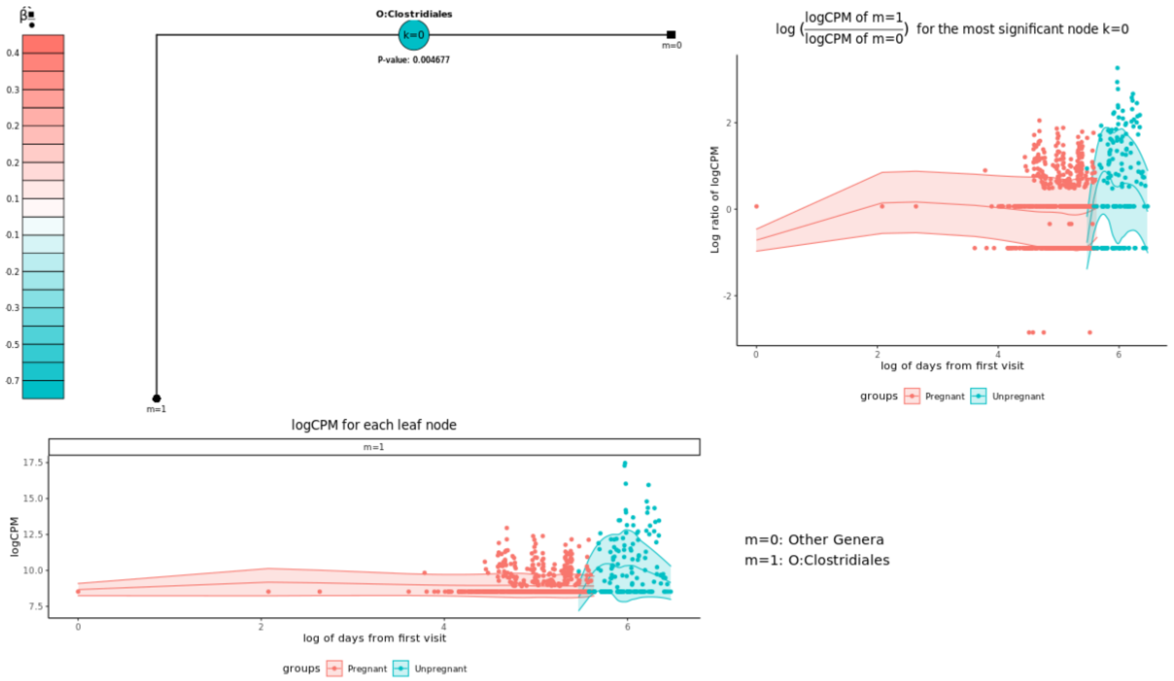


**Supplementary Figure 15. OTU distributions of significantly associated unclassified *Clostridiales*.** Relative proportions of OTUs belonging to *Clostridiales* at different time points were plotted. Each OTU has its corresponding leaf node and leaf nodes in ■ and ● indicate that they are in $\boldsymbol{L_k}$ and $\boldsymbol{R_k}$, respectively. $\widehat{\boldsymbol{\beta}}_{\frac{\blacksquare}{\bullet}}$ indicates the mean difference of $log(C^k_{ij}/D^k_{ij})$ between pregnant and unpregnant subjects after adjusting for covariates, and the red internal node indicates that OTUs in the left test leaf nodes are more abundant in pregnant subjects. The most significant node is enlarged.

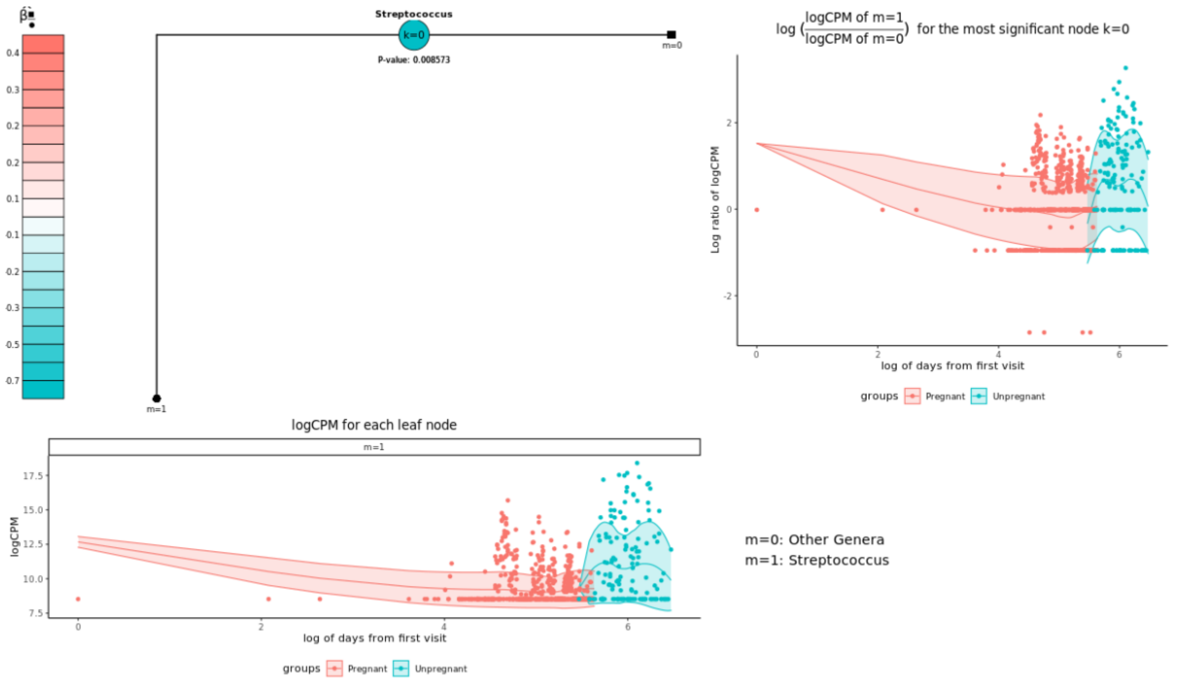


**Supplementary Figure 16. OTU distributions of significantly associated genus *Streptococcus*.** Relative proportions of OTUs belonging to *Streptococcus* at different time points were plotted. Each OTU has its corresponding leaf node and leaf nodes in ■ and ● indicate that they are in $\boldsymbol{L_k}$ and $\boldsymbol{R_k}$, respectively. $\widehat{\boldsymbol{\beta}}_{\frac{\blacksquare}{\bullet}}$ indicates the mean difference of $log(C^k_{ij}/D^k_{ij})$ between pregnant and unpregnant subjects after adjusting for covariates, and the red internal node indicates that OTUs in the left test leaf nodes are more abundant in pregnant subjects. The most significant node is enlarged.

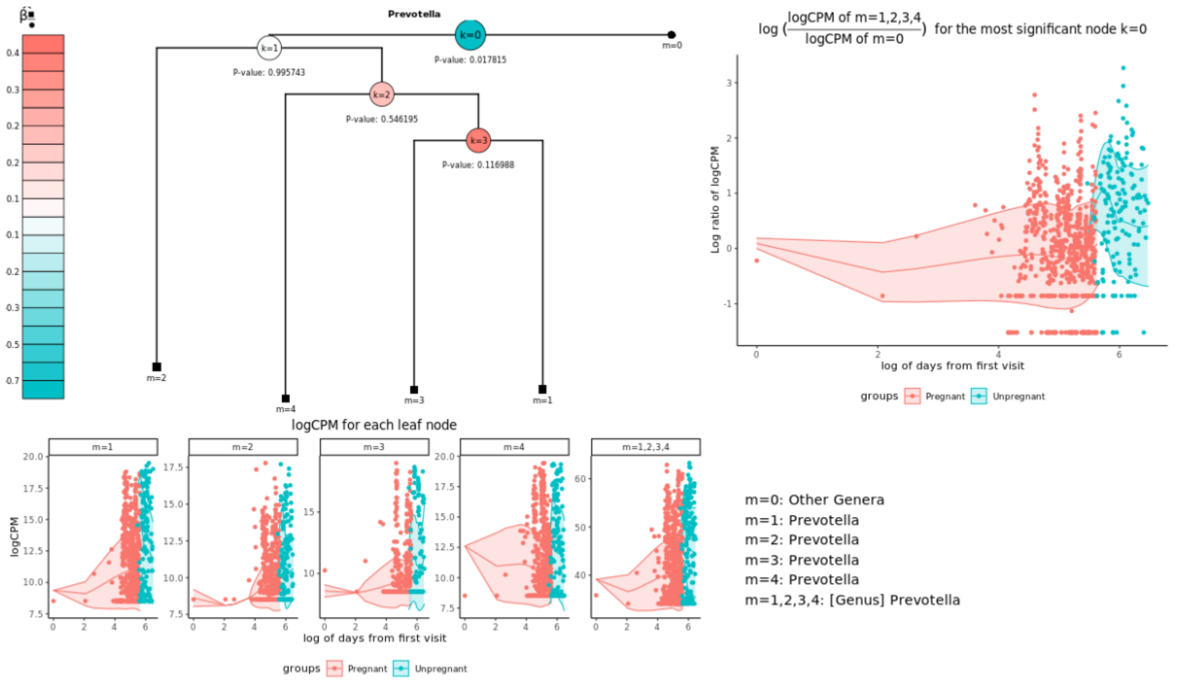


**Supplementary 17. OTU distributions of significantly associated genus *Prevotella*.** Relative proportions of OTUs belonging to *Prevotella* at different time points were plotted. Each OTU has its corresponding leaf node and leaf nodes in ■ and ● indicate that they are in $\boldsymbol{L_k}$ and $\boldsymbol{R_k}$, respectively. $\widehat{\boldsymbol{\beta}}_{\frac{\blacksquare}{\bullet}}$ indicates the mean difference of $log(C^k_{ij}/D^k_{ij})$ between pregnant and unpregnant subjects after adjusting for covariates, and the red internal node indicates that OTUs in the left test leaf nodes are more abundant in pregnant subjects. The most significant node is enlarged.
